# Specific cPRC1 complexes are co-opted to mediate oncogenic gene repression in diffuse midline glioma

**DOI:** 10.1101/2023.12.02.569451

**Authors:** Dáire Gannon, Eimear Lagan, Ademar Jesus Silva, Peter Bibawi, Anthony M. Doherty, Darragh Nimmo, Rachel McCole, Craig Monger, Giovani Genesi, Aurelie Vanderlinden Dibekeme, James A. Innes, Lu Yang, Bryan Chen, Guido van Mierlo, Pascal W.T.C Jansen, Keiran Wynne, Fran-cisco J. Sánchez-Rivera, Yadira M. Soto-Feliciano, Michiel Vermeulen, Giorgio Oliverio, Chun-Wei Chen, Richard E. Philips, Adrian P. Brackenand, Gerard L. Brien

**Affiliations:** Smurfit Institute of Genetics, Trinity College Dublin, Dublin 2, Ireland; Cancer Research UK Edinburgh Centre, Institute of Genetics and Cancer University _of Edinburgh, Edinburgh, United Kingdom; MRC Human Genetics Unit, Institute of Genetics and Cancer, The University of Edinburgh, Edinburgh, United Kingdom; Department of Neurology, Perelman School of Medicine, University of Pennsylvania, Philadelphia, Pennsylvania 19104, USA; Epigenetics Program, Perelman School of Medicine, Philadelphia, Pennsylvania 19104, USA; Abramson Cancer Center, Perelman School of Medicine, Philadelphia, Pennsylvania 19104, USA; Department of Systems Biology, Beckman Research Institute, City of Hope, Duarte, CA 91010, USA; Department of Molecular Biology, Faculty of Science, Radboud Institute for Molecular Life Sciences, Oncode Institute, Radboud University Nijmegen, 6525 GA Nijmegen, the Netherlands; Division of Molecular Genetics, The Netherlands Cancer Institute, 1066CX Amsterdam, The Netherlands; Systems Biology Ireland, University College Dublin, Belfield, Dublin 4, Ireland; Koch Institute for Integrative Cancer Research, Massachusetts Institute of Technology, Boston, Massachusetts, USA; Department of Biology, Massachusetts Institute of Technology, Cambridge, Massachusetts, USA

**Keywords:** H3K27M, Diffuse Midline Glioma, cPRC1, CBX4, PCGF4, H3K27me3

## Abstract

Diffuse midline glioma (DMG) is a fatal childhood brain tumour characterised primarily by mutant histone H3 (H3K27M). H3K27M causes a global reduction in Polycomb Repressive Complex 2 (PRC2)-mediated H3K27me3 by inhibiting PRC2 enzymatic activity. Paradoxically, PRC2 is essential in DMG tumour cells where residual complex activity is required for oncogenic gene repression, although the molecular mechanisms acting downstream of PRC2 in this context are poorly understood. Here, we’ve discovered this oncogenic gene repression is mediated by specific canonical PRC1 (cPRC1) formations. By combining CRISPR screening, biochemical and chromatin mapping approaches with functional perturbations we show that cPRC1 complexes containing CBX4 and PCGF4 drive oncogenic gene repression downstream of H3K27me3 in DMG cells. Remarkably, the altered H3K27me3 modification landscape characteristic of these tumours rewires the distribution of cPRC1 complexes on chromatin. CBX4 and PCGF4 containing cPRC1 accumulate at sites of H3K27me3 while other cPRC1 formations are displaced. Despite accounting for <5% of cPRC1 complexes in DMG, CBX4/PCGF4-containing complexes predominate as gene repressors. Our findings link the altered distribution of H3K27me3 with imbalanced cPRC1 function, promoting oncogenic gene repression in DMG cells, revealing new disease mechanisms and highlighting potential therapeutic opportunities in this incurable childhood brain tumour.

## INTRODUCTION

Diffuse Midline Gliomas (DMG) are tumours that most commonly occur in children and young adults (Mackay et al. 2017). Clinical outcomes for patients are dismal, with a median survival of less than 1 year following initial diagnosis (Jovanovich et al. 2023). These tumours can arise across several anatomical locations within the central nervous system primarily in the pons, thalamus and spinal cord (Jones and Baker 2014; Louis et al. 2016). The underlying pathological genomic features of tumours arising at different locations and ages can differ, with tumour cells activating multiple growth factor signalling pathways (Fontebasso et al. 2014; Mackay et al. 2017). However, the unifying molecular feature across all DMG tumours, regardless of anatomical site or age of onset is a chromatin signature with very low trimethylation levels at histone H3 lysine 27 (H3K27me3) (Bender et al. 2013; Lewis et al. 2013). Remarkably, this invariant molecular feature can result through multiple, non-overlapping disease mechanisms. Most tumours (∼85%) have a single, heterozygous point mutation in a histone H3 protein, which can occur in genes encoding either canonical (H3.1 and H3.2) or non-canonical (H3.3) H3 variants (Khuong-Quang et al. 2012; Schwartzentruber et al. 2012; Wu et al. 2012; Nikbakht et al. 2016). These mutations result in a lysine-to-methionine substitution at position 27 of the histone H3 tail (H3K27M). The remaining ∼15% of tumours that lack a H3K27M mutation, activate aberrantly high-level expression of the EZHIP gene (also known as CATACOMB) (Castel et al. 2020). The EZHIP protein contains a short, conversed peptide sequence that mimics the mutant H3K27M histone tail(Jain et al. 2019; Piunti et al. 2019; Jain et al. 2020). Remarkably, this peptide sequence either in the context of H3K27M or EZHIP inhibits the enzymatic activity of Polycomb Repressive Complex 2 (PRC2), which is responsible for depositing all of mono-, di- and tri-methylation at H3K27 (H3K27me1/2/3) (Conway et al. 2015). Therefore, the presence of H3K27M or EZHIP in DMG cells drives this characteristic chromatin signature.

The fact DMG tumours exploit H3K27M and EZHIP as independent means to inhibit PRC2 activity highlights the central importance of altering the H3K27 methylation landscape to tumour development. Initially, it was thought these losses would lead to the aberrant activation of PRC2 target genes by diminishing gene repressive mechanisms at these sites (Bender et al. 2013). However, we and others have shown that these losses primarily occur across broad genomic domains where the modification is transiently deposited; while H3K27me3 levels remain high at discrete genomic regions where PRC2 is more stably bound (Harutyunyan et al. 2019; Jain et al. 2020; Brien et al. 2021). We and others have established that residual PRC2 activity is essential in DMG, indicating that the discrete H3K27me3 domains that persist in tumour cells support disease biology (Mohammad et al. 2017; Piunti et al. 2017; Brien et al. 2021). Indeed, hundreds of PRC2 target genes are aberrantly repressed in H3K27M mutant cells, which depends upon the enzymatic activity of the complex (Brien et al. 2021). Therefore, a picture has emerged where the losses of H3K27me3 do not drive tumour development; but rather that retention of residual H3K27me3 at specific sites directs oncogenic gene silencing. This suggests that mechanisms downstream of H3K27me3 are important in supporting oncogenesis in this context. However, the functional impact and downstream disease mechanisms associated with these changes are largely unexplored. This limits our understanding of DMG biology and our ability to design effective therapies to treat patients.

The key functional effectors downstream of H3K27me3 are thought to be canonical forms of Polycomb Repressive Complex 1 (cPRC1) (Bracken et al. 2019; Blackledge and Klose 2021). cPRC1 complexes are biochemically defined by one of several, mutually exclusive chromatin reading CBX subunits (CBX2, CBX4, CBX6, CBX7 and CBX8) which bind PRC2 mediated H3K27me3 (Min et al. 2003; Bernstein et al. 2006; Gao et al. 2012). This is thought to be a key recruitment mechanism for cPRC1 complexes, which can mediate chromatin compaction downstream of PRC2/H3K27me3 (Francis et al. 2004; Wang et al. 2004). Thus, the combined biochemical activities of PRC2 and cPRC1 are thought to converge and mediate stable transcriptional repression (Blackledge and Klose 2021). Whether the global changes of H3K27me3 in DMG impact cPRC1 binding and/or function has not been thoroughly addressed. Moreover, whether cPRC1 function is required for the aberrant repression of PRC2 target genes in DMG is unclear. Intriguingly, emerging evidence suggests that cPRC1 function is important in DMG cells. The cPRC1 member PCGF4 (also known as BMI1) is expressed at high levels in DMG tumour cells and is required for their continued growth (Kumar et al. 2017; Balakrishnan et al. 2020; Panditharatna et al. 2022). However, the specific role(s) of cPRC1 complexes has yet to be explored in this context. In particular, the biochemical heterogeneity among discrete cPRC1 formations, defined by the distinct H3K27me3 reader CBX proteins could provide important, disease relevant functional diversity. As such, examining mechanisms downstream of H3K27me3 in DMG remains important, and will most likely provide important new insights into underlying disease biology.

Here, by combining custom and genome-wide CRISPR screening platforms, endogenous complex proteomics with quantitative chromatin mapping and functional perturbation studies we show that cPRC1 complexes containing CBX4 and PCGF4 are functionally essential in DMG cells. Significantly, this is true even though CBX4-cPRC1 complexes account for <5% of cPRC1 complexes in DMG; and while other more abundant cPRC1 formations remain functionally dispensable. Remarkably, the presence of H3K27M leads to a redistribution of cPRC1 binding on chromatin; such that CBX4 and PCGF4 containing cPRC1 complexes accumulate at PRC1/2 target sites, while biochemically distinct cPRC1 complexes containing CBX8 and PCGF2 are reduced. Notably, perturbation studies further showed that CBX4 and PCGF4 are the key gene repressors acting downstream of PRC2 and H3K27me3 in DMG cells. Taken together, our findings indicate that the altered H3K27me3 landscape in DMG causes an imbalance in cPRC1 distribution and function, leading to a predominance of gene repressive activities specifically from CBX4 and PCGF4-containing complexes; demonstrating that distinct cPRC1 complexes have non-redundant roles in gene repression, with important implications for disease and normal biology.

## RESULTS

### Residual PRC2 enzymatic activity maintains oncogenic gene repression in DMG

We and others have shown that PRC2 function is essential in DMG (Mohammad et al. 2017; Piunti et al. 2017; Brien et al. 2021). Multiple distinct PRC2 formations exist in cells based on the mutually exclusive incorporation of several non-core subunits (Figure 1A) (Hauri et al. 2016; Bracken et al. 2019; van Mierlo et al. 2019; Glancy et al. 2021). Our recent work demonstrated that different PRC2 subclasses can have distinct molecular functions (Glancy et al. 2023). Here, we wanted to understand if the PRC2 dependency in DMG is driven by any specific complex subclass. To do this, we first characterised the biochemical make-up of PRC2 using endogenous immunoprecipitation coupled with mass spectrometry (IP-MS) of the core complex members EZH2 and SUZ12 (Figure 1B and S1A). This demonstrated that the two main subclasses of PRC2 – PRC2.1 and PRC2.2 – are both present in DMG cells (Figure 1B-C and S1A-B). Stoichiometric analyses indicated that PRC2.1 is more abundant, with PRC2.1 specific accessory components comprising 77-89% of signal derived from subclass-defining members in EZH2 and SUZ12 purifications, respectively (Figure S1B-C). We used this systematic, endogenous mapping of PRC2 composition to guide the development of a saturating CRISPR/Cas9 tiling library, targeting all 11 genes encoding PRC2 members identified in DMG cells (Figure 1D). This library totalled 3,599 sgRNAs and used all PAM sequences in the coding regions of the identified PRC2 members; an approach that has been applied in other contexts to identify key functional regions within proteins of interest (Shen et al. 2015; Shi et al. 2015; Yang et al. 2021). We used this library in two independent H3K27M-mutant DMG cell lines and an additional Ewing sarcoma cell line. Strikingly, only sgRNAs targeting genes encoding the enzymatic core of EZH2, SUZ12 and EED were robustly depleted in DMG cells (Figure 1E and S1D). In contrast, in Ewing sarcoma cells which have not been reported to have any functional dependency on PRC2, no components were robustly depleted (Figure S1D). Independent validation of this screen was performed using competitive, negative selection growth assays (Figure S1E). These experiments confirmed the pooled screen results, with sgRNAs targeting EZH2, SUZ12 and EED robustly depleting. While sgRNAs targeting non-core, accessory PRC2 members elicited a limited response. Interestingly, sgRNAs targeting the additional PRC2 core member, EZH1 (EZH2 paralog) were not depleted in these experiments (Figure 1E and S1D-E). Of note, EZH1 containing PRC2 is less enzymatically active than complexes with EZH2; and EZH1 is largely dispensable for maintaining H3K27me3 in cells (Margueron et al. 2008; Shen et al. 2008). As such, only PRC2 core components that contribute significantly to enzymatic output are functionally required in DMG; indicating the maintenance of residual H3K27me3 levels in H3K27M-mutant cells underpins the PRC2 dependency in this disease context.

**Figure 1:**
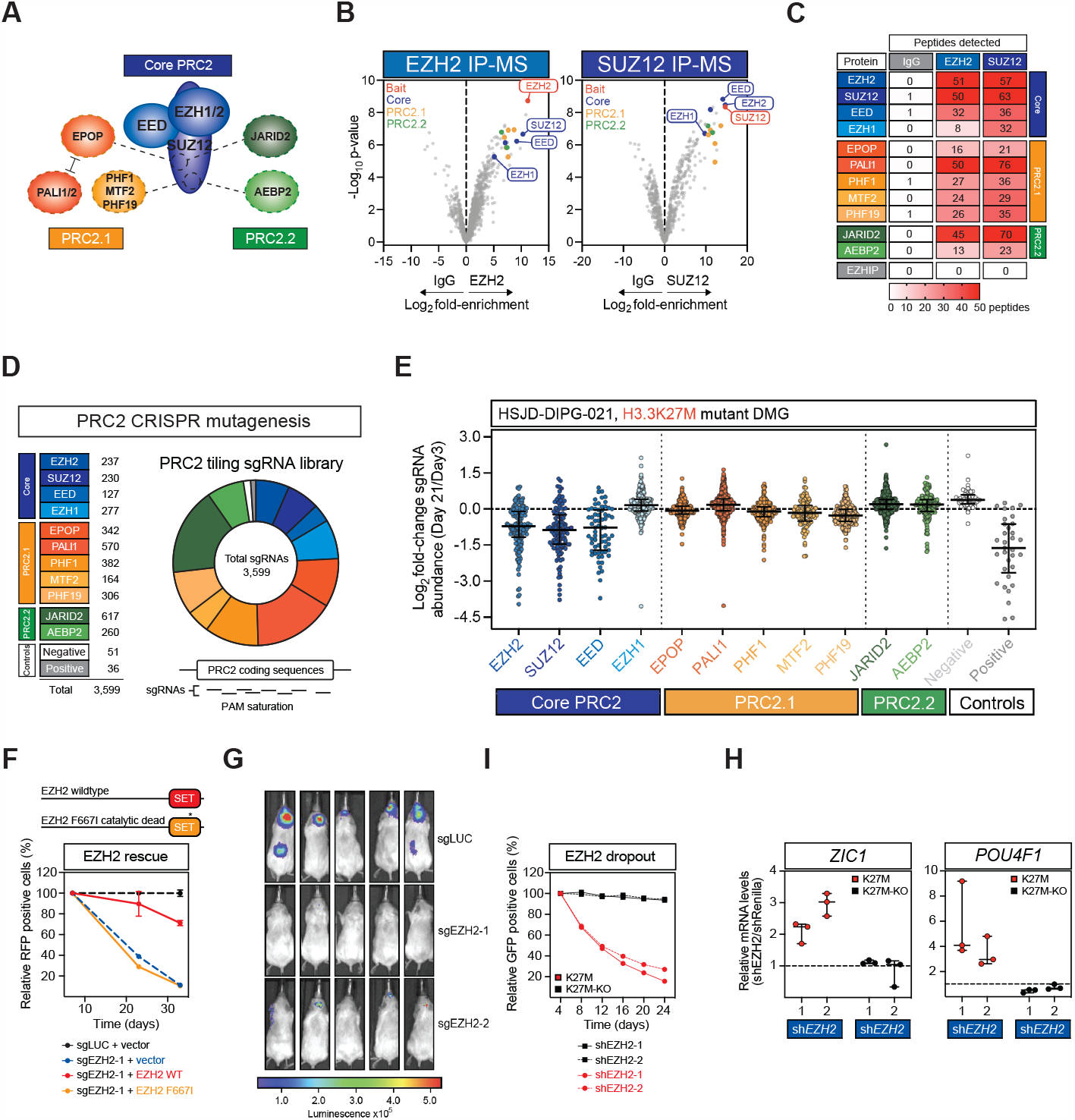
Residual PRC2 core activity maintains oncogenic gene repression in H3K27M-mutant Diffuse Midline Glioma. **A**.Schematic representation of the PRC2 core complex (blue) with PRC2.1 (orange) and PRC2.2 (green) subclass defining members. Dashed lines depict competitive binding between subclass-defining members with core component SUZ12. **B**.Volcano plots depicting protein enrichment in endogenous immunoprecipitations coupled with mass spectrometry of EZH2 (left) and SUZ12 (right) in HSJD-DIPG-007 cells. Indicated are the bait protein(s) (red), identified PRC2 core components (blue) in addition to PRC2.1 and PRC2.2 defining members (orange and green). **C**.Table depicting the number of peptides identified for each of the indicated PRC2 members in control IgG, EZH2 or SUZ12 purifications as per panel B. **D**.Schematic representation of the custom PRC2 scanning CRISPR library, indicated are the total number and proportion of sgRNAs targeting each complex member. **E**.PRC2 tiling CRISPR dropout screen in HSJD-DIPG-021 cells. Each dot represents the log2 fold-change (mean of n = 2) in the abundance of individual sgRNAs targeting the indicated PRC2 members, and controls in pooled CRISPR dropout experiments in HSJD-DIPG-021 cells. The median and interquartile range of all sgRNAs targeting a given gene are indicated. **F**.Growth competition assays in an EZH2 SET-domain functional rescue. The relative RFP+ (sgRNA+) percentage is depicted at the indicated time points after transduction. **G**.In vivo growth imaged using luciferase of EZH2 targeting and control sgRNA transduced DIPG-VI cells injected into the pons of NSG mice. **H**.Growth competition assays in DIPGXIII-K27M (red) and KO (black) cells transduced with 2 independent EZH2 targeting shRNAs. **I**.Quantitative qRT-PCRs of the indicated transcripts in DIPGXIII-K27M (red) or KO (black) cells transduced with 2 independent EZH2 targeting shRNAs (n = 3, data are represented as mean ± SD).

To further test this we performed functional rescue experiments using wildtype (WT) EZH2 or an enzymatically dead (F667I) EZH2 SET-domain mutant (Figure 1F). EZH2-WT rescued the depletion of cells transduced with a sgRNA targeting EZH2 in competitive growth assays, whereas EZH2-F667I did not (Figure 1F). To test the importance of EZH2 activity in vivo we performed xenograft experiments by injecting DMG cells transduced with control (sgLUC) or two independent EZH2 targeting sgRNAs into the pons of recipient NSG mice (Figure 1G). Disruption of EZH2 reduced overall tumour size in mice and extended survival (Figure 1G and S1F). This clearly demonstrates that EZH2/ PRC2 function is essential for robust tumorigenicity in vivo. Next, to understand if the dependency on PRC2 activity relates specifically to the presence of H3K27M we conducted orthogonal growth competition assays using EZH2 targeting shRNAs. These experiments were performed in isogenic DIPGXIII-K27M (H3K27M-positive) or DIPGXIII-KO (H3K27M-negative) DMG cell lines. Strikingly, while two independent shRNAs targeting EZH2 were strongly depleted in DIPXIII-K27M cells, they had no effect in DIPGXIII-KO cells (Figure 1I). To further understand the specific impact of EZH2 perturbation we examined the expression of PRC2 target genes, ZIC1, POU4F1, INK4A and TLE2 in this isogenic DMG cell model. Consistent with the specific growth effects, these genes were activated only in DIPGXIII-K27M, and not DIPGXIII-KO cells following EZH2 knockdown (Figure 1H and S1G). Taken together these data demonstrate PRC2 core activity is instrumental to maintain oncogenic repression of Polycomb-target genes in H3K27M-mutant cells (Brien et al. 2021; Krug et al. 2021).

### Specific cPRC1 complex formations are functionally essential in DMG

To identify a more comprehensive list of genetic dependencies in DMG we performed additional genome-wide CRISPR/Cas9 functional screening experiments in the same DMG cell lines used in our PRC2 focussed scanning screens (Figure 1). This again revealed that only the core PRC2 members EZH2, SUZ12 and EED scored as strong dependencies, whereas non-enzymatic PRC2 accessory components were dispensable (Figure 2A-B and S2A). However strikingly, two specific members of cPRC1 complexes – CBX4 and PCGF4 – had comparable dependency scores to the PRC2 core components in these experiments (Figure 2C and S2A). Both CBX4 and PCGF4 are part of larger paralog groups, which assemble into mutually exclusive cPRC1 formations (Figure 3D) (Gao et al. 2012). Surprisingly, no other genes encoding cPRC1 members scored as dependencies in these experiments (Figure 2C and S2A). This suggests that CBX4 and PCGF4 may execute specific, non-redundant functions that cannot be compensated for by their paralogs (Figure 2C). Independent sgRNA depletion assays validated this pooled screen result with sgRNAs targeting CBX4, but not its paralogs CBX2, CBX6, CBX7 or CBX8 robustly depleting in multiple independent DMG cell lines (Figure 2E and S2B). Moreover, sgRNAs targeting PCGF4 but not its paralog PCGF2 (also known as MEL18) were robustly depleted in these validatory experiments (Figure 2E). The specificity of the CBX4 dependency was further confirmed in a separate custom CRISPR screening experiment using a library of 2,303 sgRNAs targeting ∼550 chromatin regulatory genes (Figure S2C). This screen was conducted in an additional, independent H3K27M-mutant DMG cell line, DIPGVI, and we observed that all sgRNAs targeting CBX4 were significantly depleted, while no sgRNAs targeting CBX2, CBX6, CBX7 or CBX8 had any effect (Figure S2C). In an orthogonal shRNA depletion assay using an independent H3K27M-mutant DMG cell line, we saw specific depletion of shRNAs targeting CBX4 and PCGF4, but not their respective paralogs (Figure S2D). Previous reports indicate that CBX4 can function as an E3-SUMO ligase, independent of cPRC1 (Kagey et al. 2003; Roscic et al. 2006; Li et al. 2007; Li et al. 2014). To understand, if CBX4 may be functioning independently of cPRC1 in DMG we performed endogenous IP-MS to identify CBX4 associated proteins. Although CBX4 did co-purify proteins not part of cPRC1 (Figure S2E); gene ontology analysis of these proteins showed a stark enrichment of PRC1 and terms related to chromatin regulation (Figure S2F). Moreover, none of the SUMOylation related proteins previously linked to CBX4 could be identified in these purifications. Together, these data indicate this dependency is related to CBX4 in the context of cPRC1 and the functional role of these specific complexes in DMG tumour cells.

**Figure 2:**
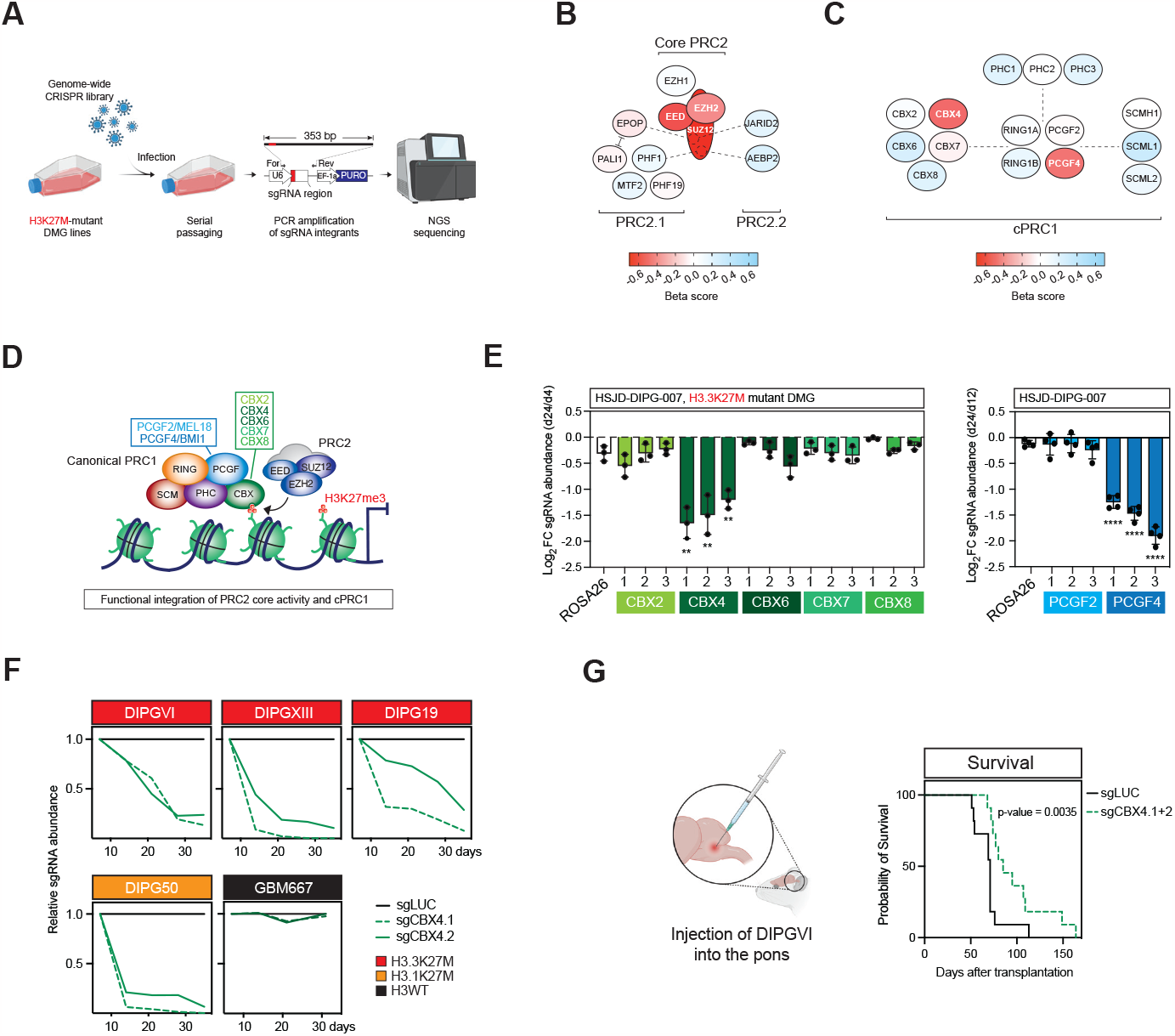
CBX4 and PCGF4 containing cPRC1 complexes are functionally essential in DMG. **A**.Schematic overview of the genome-wide CRISPR functional screening approach in DMG cells **B**.Heatmap representation of CRISPR dependency scores for all PRC2 members from screening experiments in genome-wide screens in HSJD-DIPG-021. **C**.Heatmap representation of CRISPR dependency scores for all cPRC1 members from screening experiments in genome-wide screens in HSJD-DIPG-021. **D**.Schematic representation illustrating the functional integration of cPRC1 complexes and PRC2, via the direct binding of H3K27me3 by individual CBX paralogs. **E**.Growth competition assays in HSJD-DIPG-007-Cas9 cells expressing sgRNAs targeting each of the indicated CBX (left) and PCGF (right) genes (n = 3 or 4, data represents mean ¬± SD). P-values calculated using Student’s t-test comparing against negative control sgRNA targeting ROSA26, * = P ≤ 0.05, ** = P ≤ 0.01, *** = P ≤ 0.001 and **** = P ≤ 0.0001. **F**.Growth competition assays in the indicated DMG and glioma cell lines expressing two independent CBX4 targeting or a control (sgLUC) sgRNA. The histone H3 genotype of each cell line is colour-coded and indicated on the right. **G**.Survival curve for recipient mice injected with DIPGVI cells expressing control (sgLUC) or two independent CBX4 targeting sgRNAs. Represented is the survival difference between mice injected with sgLUC cells and the average of the two independent CBX4 targeted conditions.

**Figure 3:**
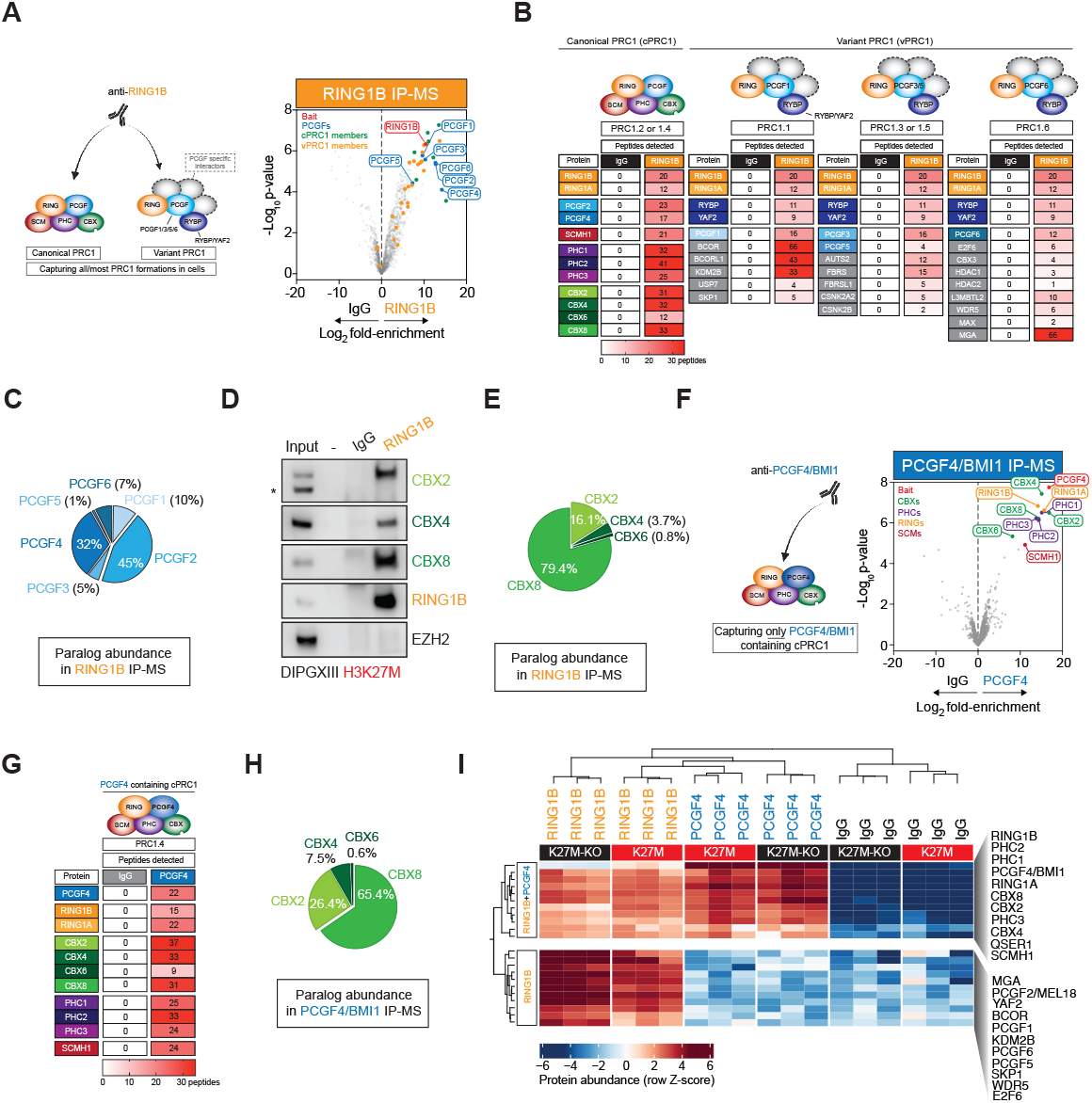
CBX4 and PCGF4 containing complexes account for a minor fraction of cPRC1 in DMG. **A**.Schematic representation of RING1B IP-MS approach capturing PRC1 subclasses (Left), with a volcano plot of endogenous RING1B IP-MS in HS-JD-DIPG-007. cPRC1, vPRC1, PCGF paralogs and RING1B are colour-coded and indicated on the plot. **B**.Table depicting peptide numbers identified for each of the indicated PRC1 members in RING1B IP-MS in HSJD-DIPG-007, with the table separated based on PRC1 molecular subclass. **C**.Pie chart denoting the percent of total PCGF protein abundance accounted for by each of the individual PCGF paralogs in RING1B IP-MS in HS-JD-DIPG-007. **D**.Immunoblots of the indicated PRC1/2 proteins in RING1B IPs in DIPGXIII cells. **E**.Pie chart denoting the percent of total CBX protein abundance accounted for by each of the individual CBX paralogs in RING1B IP-MS in HSJD-DIPG-007. **F**.Schematic representation of PCGF4 IP-MS approach (left), with a volcano plot of endogenous PCGF4 IP-MS in DIPGXIII-K27M. All cPRC1 members identified are indicated. **G**.Table depicting peptide numbers identified for each of the indicated cPRC1 members in PCGF4 IP-MS in DIPGXIII-K27M. **H**.Pie chart denoting the percent of total CBX protein abundance accounted for by each of the individual CBX paralogs in PCGF4 IP-MS in DIPGXIII-K27M. **I**.Heatmap depicting enrichment of the indicated proteins in RING1B and/or PCGF4 IP-MS in DIPGXIII-K27M or KO cells.

Interestingly, additional CBX4 sgRNA depletion assays suggest that the CBX4 dependency is linked specifically to the presence of H3K27M. While we saw robust depletion of sgRNAs targeting CBX4 in multiple H3K27M-mutant DMG cell lines, in either a histone H3.1 or H3.3 context; in an additional glioma cell line with wildtype H3, CBX4 appeared dispensable (Figure 2F). Moreover, in genome-wide CRISPR screening data from 1,095 cancer cell lines representing 28 different lineages, CBX4 and PCGF4 were dependencies in <1.5% of cell lines, highlighting that they are not common essential genes (depmap.org) (Figure S2G). Consistent with our findings in tissue culture models, DMG cells transduced with CBX4 targeting sgRNAs had reduced tumour-forming capacity following injection into recipient mice (Figure 2F). Taken together, these data indicate that CBX4/PCGF4-containing cPRC1 complexes are specifically functionally essential in DMG cells; and highlight the importance of these complexes for tumorigenicity in vivo.

### CBX4 and PCGF4 containing complexes constitute a minor pool of cPRC1

To understand if the specific functional dependency on CBX4 and PCGF4-containing complexes simply stems from them being in the predominant form(s) of PRC1 in DMG, we performed endogenous IP-MS of RING1B in two independent DMG cell lines (Figure 3A and S3A). RING1B provides the broadest view of PRC1 assembly since it captures both the canonical and variant PRC1 subclasses (Gao et al. 2012; Kloet et al. 2016; Bracken et al. 2019). Analyses of these experiments found significant heterogeneity among purified PRC1 complexes (Figure 3A). In addition to PCGF2/4, which are primarily found in cPRC1 complexes; the vPRC1 specific PCGF1/3/5/6 paralogs all co-purified with RING1B (Figure 3A-B and S3A-B). Notably, cPRC1 components were more abundant than many vPRC1 members, suggesting cPRC1 complexes predominate in DMG cells (Figure 3A-B and S3A-B). Consistent with this, a closer examination of PCGF protein stoichiometry showed the majority of signal was derived from cPRC1 associated PCGF2/4 (Figure 3C and S3C). Collectively, PCGF2 and PCGF4 account for 77-85% of total PCGF protein signal in the two DMG cell lines; indicating that most assembled complexes are of the cPRC1 subclasses (Figure S3C). While vPRC1 containing either PCGF1 or PCGF6 account for most of the remaining complexes (Figure 3C). Significant compositional heterogeneity is evident within RING1B-associated cPRC1 complexes (Figure 3B and S3B). Multiple mutually exclusive cPRC1 members co-purify with RING1B, including all three PHC proteins (PHC1-3), four out of five CBXs (CBX2/4/6/8) and PCGF2/4 (Figure 3B and S3B-D). These data demonstrate that a heterogeneous mix of different PRC1 formations are present in DMG cells, with cPRC1 assemblies accounting for most complexes.

Multiple CBX paralogs co-purified with RING1B in DMG cells (Figure 3D). To understand if CBX4 is the predominant CBX present in this context, we examined CBX paralog stoichiometry within RING1B-associated complexes. Strikingly, this revealed that CBX4 accounted for only a very small portion (1.8-3.7%) of the overall CBX protein signal (Figure 3E and S3E). Remarkably, despite being functionally dispensable (Figure 2 and S2), CBX8 alone accounted for the majority of CBX protein signal (79.4-83.6%), while CBX2 accounted for most of the remaining CBX abundance (13.7-16.1%) (Figure 3E and S3E). Collectively, CBX2 and CBX8 account for over 95% of CBX signal in RING1B-containing complexes, with CBX6 comprising less than 1% in each cell line. Therefore, despite CBX4 being the essential CBX paralog in terms of function, it is present in less than 5% of RING1B-associated PRC1 complexes in DMG cells. Since PCGF4 is also a functional dependency we wondered whether it preferentially forms complexes with CBX4 in DMG cells. To test this, we performed additional endogenous IP-MS using an antibody recognising PCGF4 (Figure 3F). Similar to RING1B purifications, significant heterogeneity was evident among PCGF4-containing cPRC1 complexes (Figure 3F-G). PCGF4 co-purified both RING proteins (RING1A/B), all three PHCs (PHC1-3) and the same four CBX proteins (CBX2/4/6/8) present in RING1B-containing complexes (Figure 3F-G and S3F). Indeed, stoichiometric analysis of PCGF4 purifications demonstrated the CBX proteins exist in comparable proportions to those seen in RING1B purifications (Figure 3E, H and S3E). Therefore, PCGF4 does not preferentially associate with CBX4 over CBX2 or CBX8, which are the predominant CBX partners for PCGF4 (Figure 3H). Finally, we wanted to understand if the assembly dynamics of PRC1 complexes are affected by the presence of H3K27M. To address this, we performed additional RING1B and PCGF4 IP-MS in isogenic DIPGXIII-K27M versus DIPGXIII-KO cells (Figure 3I). This demonstrated that the overall assembly dynamics of both vPRC1 and cPRC1 are unchanged between DIPGXIII-K27M and DIPGXIII-KO cells (Figure 3I). Taken together these data demonstrate that while a heterogenous mix of cPRC1 complexes are present in DMG cells (Figure S3G), CBX4 and PCGF4 containing complexes constitute just a minor (<5%) pool of complexes in this context. Moreover, this implies that the additional CBX and PCGF paralogs (CBX2, CBX8 and PCGF2) in other cPRC1 formations are not functionally redundant with CBX4/PCGF4, suggesting despite their low abundance, complexes with CBX4/PCGF4 have specific function(s).

### PRC1 chromatin binding dynamics are altered in H3K27M-mutant cells

Next, we wanted to evaluate whether changes in cPRC1 occupancy might explain the functional dependency on CBX4 and PCGF4 containing cPRC1 complexes. To test this, we quantitatively mapped PRC1/2 members in isogenic H3K27M-mutant and WT cells (Figure 4A and S4A). We initially examined cPRC1 and PRC2 occupancy using CUT&RUN-qPCR in DIPGXIII-K27M and KO cells (Figure S4A). This showed that SUZ12, RING1B, CBX4 and PCGF4 levels all increased at a PRC1/2 target site; while CBX8 and PCGF2 levels were both reduced (Figure S4A). These changes motivated us to establish a quantitative CUT&RUN-Rx approach to accurately examine global cPRC1/PRC2 dynamics in H3K27M-mutant cells (Figure 4A). To facilitate these global CUT&RUN measurements we identified cPRC1 and PRC2 antibodies that cross-react with their cognate target in both human and mouse cells; and adapted the standard CUT&RUN protocol by adding 10% mouse cells as an exogenous spike-in to human glioma cells (Orlando et al. 2014; Skene and Henikoff 2017). Importantly, we could derive high-quality reads mapping to the spike-in genome for all antibodies (Figure S4B); allowing us to calculate quantitative spike-in normalisation factors to apply to data derived from DMG cells (see Methods). We first identified PRC1/2-bound genomic regions (4,475) based on overlapping RING1B and SUZ12 peaks (Figure S4C). As we and others have previously shown SUZ12 (PRC2) binding increased at Polycomb-bound regions in H3K27M mutant cells (Figure S4C) (Jain et al. 2020; Brien et al. 2021). Consistent with our CUT&RUN-qPCRs, global analyses of RING1B, CBX4 and PCGF4 demonstrated that complexes with these proteins were elevated at Polycomb-bound regions, while complexes with CBX8 and PCGF2 were reduced in H3K27M-mutant cells (Figure 4B-C and S4C). The use of spike-in normalisation was essential here to have an accurate global view of cPRC1 dynamics. Use of standard read depth normalisation misrepresented global changes for both PCGF2 and CBX8 (Figure S4D). Spike-in normalised data matched our quantitative CUT&RUN-qPCRs showing reduced abundance of PCGF2 and CBX8 in H3K27M-mutant cells; while read depth normalisation showed the opposite (Figure S4A&D). It has been established that use of a reference genome spike-in in chromatin mapping approaches is essential to quantitatively map global losses; where lacking such normalisation can underestimate or misrepresent changes (Orlando et al. 2014). Importantly, these dynamics were recapitulated for CBX4 and CBX8 in an independent mouse NSC model, exogenously expressing WT or K27M-mutant histone H3.3 (Figure S4E). This demonstrates that these opposing dynamics, with both increases and decreases for different cPRC1 member paralogs, is a general feature of H3K27M-mutant cells.

**Figure 4:**
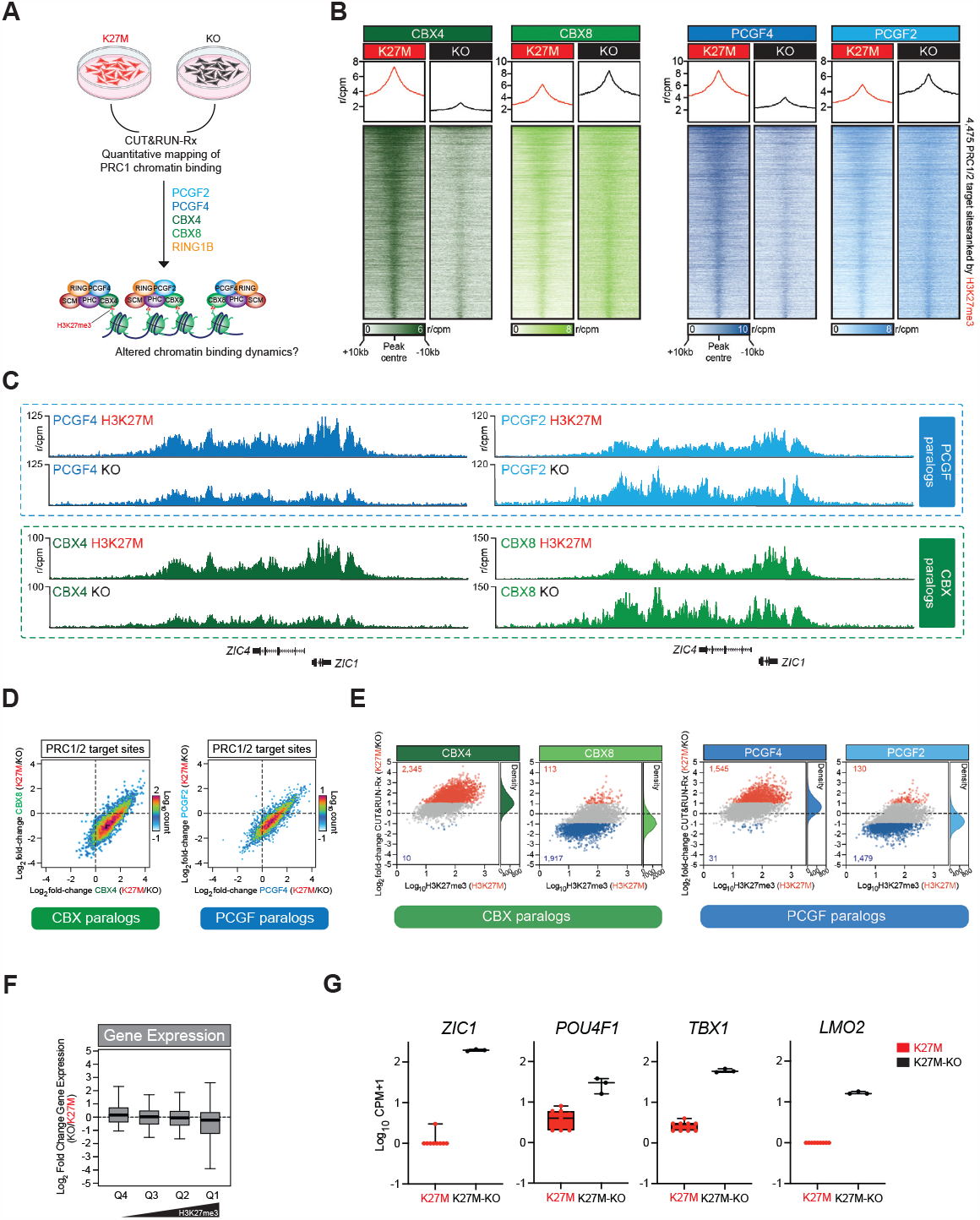
Specific accumulation of CBX4 and PCGF4 at Polycomb target sites in DMG cell. **A**.Schematic representation of the quantitative CUT&RUN approach mapping multiple cPRC1 members in DIPGXIII-K27M and KO cells. **B**.Tornado and average plots showing average enrichments of cPRC1 members in biological triplicate CUT&RUN-Rx mapping in DIPGXIII-K27M and KO cells. Indicated are the regions corresponding to the genomic window +/-10kb of PRC1/2 target sites. The scale bar denotes reference-adjusted counts per million mapped reads (r/CPM) for all experiments. **C**.Genomic tracks showing average quantitative PCGF4, PCGF2, CBX4 and CBX8 CUT&RUN-Rx signal from biological triplicate experiments at the indicated genomic locus in DIPGXIII-K27M and KO cells. **D**.XY scatter plot correlating changes in CBX4 and CBX8 (left) or PCGF2 and PCGF4 (right) binding at PRC1/2 target sites in DIPGXIII-K27M and KO cells. **E**.MA-plots showing fold-change in binding for CBX4, CBX8, PCGF4 and PCGF2 in at PRC1/2 target sites in DIPGXIII-K27M and KO cells. Sites changing >2-fold are shown in red (increasing) or blue (decreasing) with the number of sites in these categories also indicated. The densities of chromatin binding changes are shown on the right of each plot. **F**.Boxplots showing log2 fold change in RNA abundance of PRC1/2 target genes between DIPGXIII-K27M and KO cells; genes are separated into quartiles based on their local enrichment of H3K27me3 in DIPGXIII-K27M cells.

We next wanted to understand if these changes represented a competitive relationship between the CBX and PCGF paralogs where increases in one, came at the expense of the other. However, comparing the binding dynamics of CBX4 versus CBX8, and PCGF2 versus PCGF4 showed these dynamics were not (anti)-correlated (Figure 4D). In fact, sites with the greatest increases in CBX4 and PCGF4 binding had relatively stable CBX8 and PCGF2 levels (Figure 4D); consistent with an overall increase in PRC1 abundance at these sites (Figure S4C). Interestingly, underlying levels of H3K27me3 appear to dictate these cPRC1 dynamics (Figure 4E and S4F). Sites with the highest levels of H3K27me3 had the greatest increase in CBX4 and PCGF4 while retaining near-normal (slightly reduced) levels of CBX8 and PCGF2 (Figure 4E and S4F-G). Consistent with the idea underlying H3K27me3 levels are instrumental in the redistribution of cPRC1, quantitative mapping of the vPRC1-specific protein RYBP demonstrated that although overall levels slightly increased at Polycomb-bound sites (Figure S4H); these increases did not have any association with underlying H3K27me3. Taken together, these data demonstrate that cPRC1 binding dynamics are altered in H3K27M-mutant cells, leading to a specific increase in complexes containing CBX4 and PCGF4 at PRC1/2 bound genomic regions.

Next, we examined gene expression dynamics of Polycomb target genes in DIPGXIII-K27M versus KO cells (Figure 4F and S4I). We have previously shown that a subset of strong Polycomb target genes have reduced expression levels in H3K27M-mutant human NSCs, and that this is associated with increased PRC2 binding at their promoters (Brien et al. 2021). Here, we separated Polycomb targets into quartiles based on H3K27me3 levels and found that genes with the highest levels of the modification have reduced gene expression in DIPGXIII-K27M cells (Figure 4F-G and S4J). Therefore, consistent with our previous findings in an independent DMG model system, many Polycomb target genes are aberrantly repressed in H3K27M-mutant cells. Notably, these genes are also the sites where CBX4/PCGF4 binding increase the most (Figure 4E and S4G), suggesting that changes in cPRC1 distribution, and accumulation of CBX4/PCGF4 containing complexes may drive this aberrant silencing.

### CBX4 and PCGF4 drive oncogenic repression of Polycomb targets

We previously demonstrated that the aberrant repression of Polycomb target genes in H3K27M-mutant cells could be reversed by PRC2 inhibition (Brien et al. 2021). Given the specific functional importance of CBX4 and PCGF4, we wondered whether PRC2 activity might preferentially support the function of CBX4/PCGF4-containing cPRC1 in DMG cells. To test this, we treated DIPGXIII-K27M cells with the EZH2 inhibitor Tazemetostat. Over a treatment time course of 3, 6 and 9 days we observed an accumulation of gene expression changes (Figure 5A). Consistent with previous observations that EZH2 inhibition requires 5-7 days to elicit a robust phenotypic impact (Knutson et al. 2013; Knutson et al. 2014), at day 3, just 23 genes were differentially expressed all of which were upregulated (Figure 5A). However, consistent with the on-target activity of Tazemetostat, 18 (78%) of these genes were direct Polycomb targets (Figure S5A). Following 6 and 9 days of treatment, the number of differentially expressed genes accumulated to 465 and 2,082, respectively (Figure 5A). Gene ontology (GO) analyses demonstrated a progressive enrichment of neurodevelopmental terms among upregulated genes; and DNA replication and cell cycle among those downregulated (Figure S5B). This supports previous findings that PRC2 inhibition promotes both the differentiation and cell cycle arrest of DMG cells (Brien et al. 2021). We decided to map cPRC1 binding dynamics after 6 days of Tazemetostat treatment since many upregulated genes 145 of 316 (46%) – were direct Polycomb targets (Figure 5A and S5A). In contrast, the vast majority of differentially expressed genes by day 9 were not Polycomb targets, likely reflecting the accumulation of secondary downstream changes (Figure S5A). Quantitative CUT&RUN-Rx mapping of CBX4, CBX8, PCGF2 and PCGF4 showed that all cPRC1 members were displaced to a comparable level following PRC2 inhibition and the associated loss of H3K27me3 (Figure 5B-C and S5C). This suggests that PRC2 and H3K27me3 do not preferentially support the chromatin binding of any particular cPRC1 assembly. Indeed, the loss of each CBX and PCGF paralog was closely correlated at Polycomb-bound sites (Figure 5D); further supporting the idea these complexes are similarly dependent on PRC2 activity for their chromatin binding. We recently showed the physical presence of JARID2 containing PRC2.2 can support the chromatin binding of cPRC1 complexes containing CBX7 in embryonic stem cells (Glancy et al. 2023). Although CBX7-cPRC1 is not present in DMG cells (Figure 3); this motivated us to quantitatively map PRC2 (SUZ12) in Tazemetostat-treated DMG cells (Figure S5D). This showed PRC2 is also displaced from many Polycomb-bound sites, consistent with potential H3K27me3 dependent- and independent contributions of PRC2 to cPRC1 recruitment being disrupted by Tazemetostat treatment (Figure S5D). Interestingly, while cPRC1 members were depleted from essentially all Polycomb-bound loci (Figure 5B-D); SUZ12 levels were unchanged at a subset of regions (Figure S5E-F). Sites which retained SUZ12 were typically regions with high levels of H3K27me3 in untreated cells, indicating that Tazemetostat treatment and loss of H3K27me3 does not disrupt PRC2 recruitment at many of the strongest Polycomb-target sites (Figure S5F). In contrast, cPRC1 members exhibited the greatest loss from these strong Polycomb-target sites (Figure S5F), suggesting that the physical presence of PRC2 is not a dominant recruitment mechanism in these regions. Consistent with the idea H3K27me3 predominates in this regard, loss of cPRC1 tracks with H3K27me3 across all Polycomb-target sites (Figure 5E and S5F-G). The sites with the greatest reduction of H3K27me3 have the greatest displacement of cPRC1 members (Figure 5E). This reduced cPRC1 binding was associated with activation of Polycomb-target genes, with progressively increasing levels of gene expression apparent with greater cPRC1 displacement (Figure 5E). Notably, the sites where CBX4 and PCGF4 accumulated the most in H3K27M-mutant cells (Figure 4F and S4E); were the same as those losing cPRC1 members and activating gene expression to the greatest degree following Tazemetostat treatment (Figure 5E). Taken together, this suggests that increased CBX4/ PCGF4 binding at these sites may be responsible for their aberrant repression in H3K27M-mutant cells; and that the effects of Tazemetostat are mediated specifically via displacement of CBX4/PCGF4.

**Figure 5:**
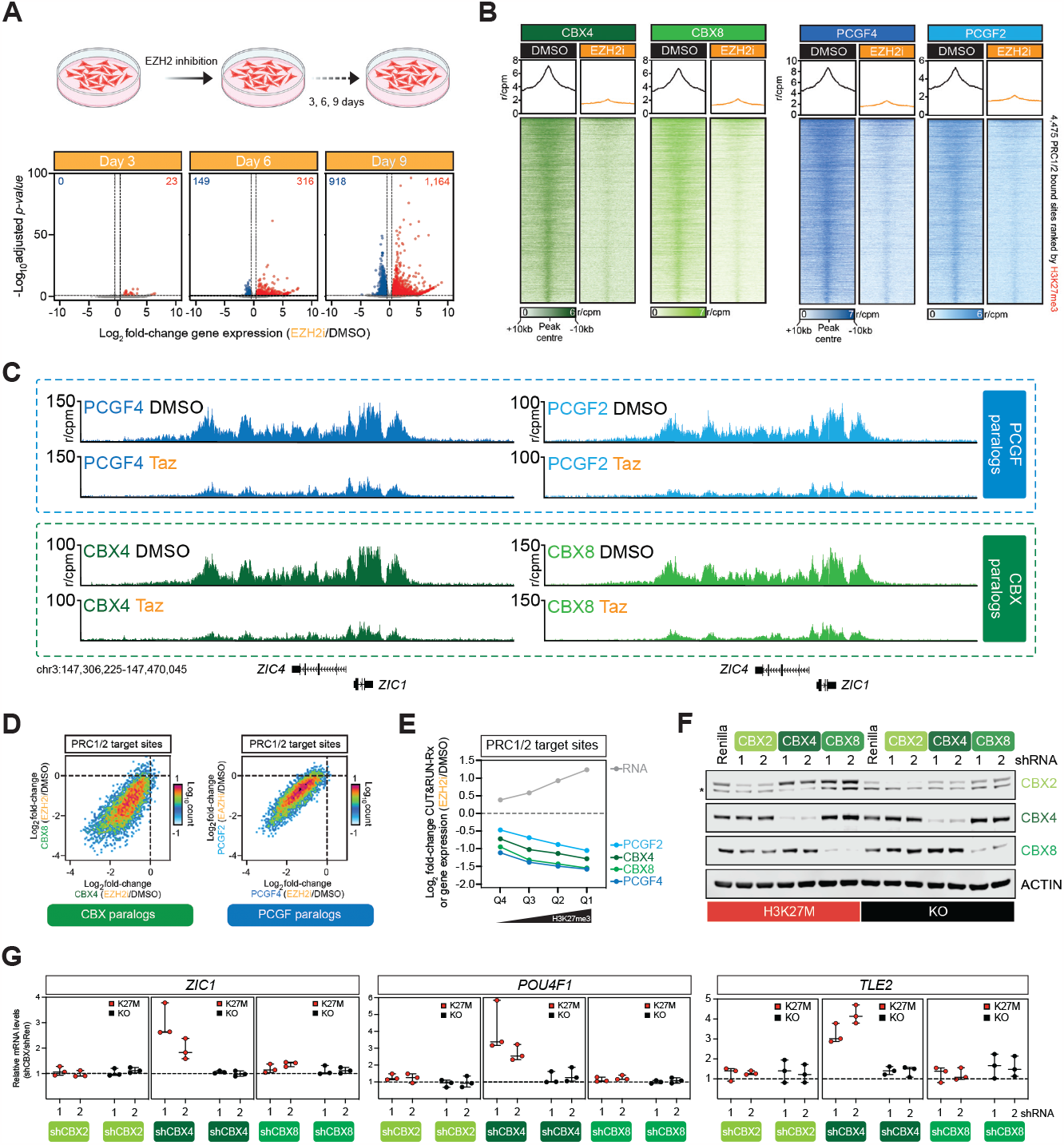
CBX4 and PCGF4 containing cPRC1 complexes specifically drive oncogenic gene repression downstream of H3K27me3 in DMG. **A**.Schematic showing EZH2 inhibitor treatment time course in DIPGXIII-K27M cells (Top); with volcano plots presenting differentially expressed genes at days 3, 6 and 9 following EZH2 inhibitor treatment at a concentration of 2□M. The number of genes differentially expressed both up (red) and down (blue) are indicated at each time point. **B**.Tornado and average plots showing average enrichments of cPRC1 members in biological triplicate CUT&RUN-Rx mapping in DIPGXIII-K27M cells treated with EZH2 inhibitor for 6 days. Indicated are the regions corresponding to the genomic window +/-10kb of PRC1/2 target sites. The scale bar denotes reference-adjusted counts per million mapped reads (r/CPM) for all experiments. **C**.Genomic tracks showing average quantitative PCGF4, PCGF2, CBX4 and CBX8 CUT&RUN-Rx signal from biological triplicate experiments at the indicated genomic locus in control (DMSO) treated DIPGXIII-K27M or cells treated with EZH2 inhibitor for 6 days. **D**.XY scatter plots correlating changes in CBX4 and CBX8 (left) or PCGF2 and PCGF4 (right) binding at PRC1/2 target sites in DIPGXIII-K27M cells treated with EZH2 inhibitor for 6 days. **E**.Dynamics of chromatin and gene expression changes at PRC1/2 target gene quartiles in DIPGXIII-K27M cells treated with EZH2 inhibitor for 6 days. Shown are the mean changes of the indicated cPRC1 members and RNA levels relative to control (DMSO) treated cells. **F**.Immunoblot of the indicated proteins in DIPGXIII-K27M (red) or KO (black) cells expressing control Luciferase or 2 independent shRNAs targeting CBX2/ CBX4/CBX8. *Denotes a background protein in CBX2 blots. **G**.Quantitative qRT-PCRs of the indicated transcripts in DIPGXIII-K27M (red) or KO (black) cells transduced with 2 independent shRNAs targeting CBX2, CBX4 and CBX8 (n = 3, data represents mean ¬± SD).

To evaluate if there were any key functional contributions from specific cPRC1 complexes relating to aberrant Polycomb-target gene repression, we used an independent shRNA-mediated knockdown approach to specifically deplete CBX2, CBX4, CBX8, PCGF2 or PCGF4 in DIPGXIII-K27M and KO cells (Figure 5F and S5H). Having established this knockdown approach, we examined the transcript levels of several Polycomb-target genes activated in Tazemetostat-treated cells. Strikingly, these genes were only activated following knockdown of CBX4 or PCGF4, but not CBX2, CBX8 or PCGF2 (Figure 5G and S5I). Since PCGF4 is the predominant PCGF paralog in DIPGXIII cells, it is perhaps unsurprising that knockdown of PCGF4 (but not PCGF2) would activate Polycomb-target genes. However, despite accounting for <2% of total CBX paralog abundance in these cells, CBX4 predominates in mediating oncogenic gene repression (Figure 5G). Remarkably, these effects were specific to H3K27M-mutant cells since knockdown of CBX/PCGF paralogs had limited, or no effect on expression of the same Polycomb targets in isogenic DIPGXIII-KO cells (Figure 5G and S5H). This is despite the fact cPRC1 composition is unchanged between DIPGXIII-K27M and KO cells (Figure 3I). Therefore, like EZH2, CBX4 and PCGF4 are specifically required to maintain oncogenic gene repression in H3K27M-mutant cells (Figure 1H and 5G). Taken together, these findings demonstrate that specific cPRC1 complexes drive oncogenic gene repression downstream of PRC2 activity in H3K27M mutant DMG tumour cells (Figure 5H).

## DISCUSSION

Altered chromatin regulation is a pervasive hallmark of oncogenesis in childhood malignancies (Grobner et al. 2018; Hanahan 2022). Mechanistic dissection of this altered chromatin biochemistry is providing important knowledge of disease biology; but also, compelling insights into fundamental aspects of chromatin regulation (Brien et al. 2016; Bracken et al. 2019). Here, we’ve shown that the presence of the H3K27M oncohistone in DMG leads to a specific functional reliance on cPRC1 complexes containing CBX4 and PCGF4. PRC2 core complex methyltransferase activity maintains a disease associated H3K27me3 landscape, leading to the redistribution of cPRC1 complexes on the chromatin fibre. Despite accounting for less than 5% of overall cPRC1 abundance, accumulation of CBX4 and PCGF4 containing cPRC1 complexes drives oncogenic gene repression in DMG cells. These findings establish how altering the reading dynamics of H3K27me3 influences downstream gene repression, highlighting the non-overlapping gene repressive activities of distinct cPRC1 complexes. Moreover, we’ve discovered new avenues for the development of mechanistically-anchored therapeutics in this incurable disease.

Disrupted chromatin regulation has been proposed to disrupt normal developmental processes as a fundamental step in oncogenesis childhood cancers. Consistent with this, DMG tumour cells are enriched in primitive oligodendrocyte precursor cells which appear less capable of undergoing differentiation (Filbin et al. 2018; Jessa et al. 2019). H3K27M is believed to limit the differentiation potential of DMG cells due to the aberrant repression of PRC2 target genes (Brien et al. 2021; Krug et al. 2021). However, the molecular mechanisms underlying this aberrant repression have been unclear. We’ve shown that CBX4/PCGF4-containing cPRC1 complexes are central in this regard. Mechanistically, our data suggest the altered H3K27me3 landscape characteristic of these tumours drives this functional dependency. In H3K27M and EZHIP expressing DMG cells, H3K27me3 distribution changes from a relatively diffuse pattern where the modification is present throughout much of the genome, to a much more discrete pattern with H3K27me3 aberrantly focused at stable PRC2 binding sites (Harutyunyan et al. 2019; Jain et al. 2020; Brien et al. 2021). Therefore, beyond simply reducing H3K27me3 levels, H3K27M and EZHIP shift the relative distribution pattern of the modification. These changes correlate with the redistribution of CBX4 and PCGF4-containing cPRC1 complexes to H3K27me3 enriched sites; suggesting the pattern of H3K27me3 distribution differentially instructs chromatin binding dynamics of different cPRC1 complex configurations. Where altering this balance can have significant downstream functional impact(s). Interestingly, an independent childhood brain tumour – Posterior Fossa Type A (PFA) ependymoma – is characterised by a similar H3K27me3 landscape to DMG tumours (Krug et al. 2021). PFA ependymoma has a shared aetiology to DMG with the vast majority (95%) of patients having elevated EZHIP expression, and the remaining patients carrying a H3K27M mutation (Ryall et al. 2017; Pajtler et al. 2018). Genomics studies have yet to find any significantly co-occurring genetic abnormalities in these tumours, suggesting that in this context EZHIP/H3K27M are sufficient to drive oncogenesis. Strikingly, PFA tumour cells are also functionally reliant on CBX4 and PRC2 core components (Mack et al. 2014; Michealraj et al. 2020). The shared aetiology, chromatin landscape and functional dependencies in DMG and PFA is striking and supports the idea that the altered pattern of H3K27me3 distribution is linked to the functional dependency on CBX4. Molecular studies of cPRC1 in PFA ependymoma have yet to be reported, so it remains important to examine CBX4 in this context. However, given the clear links between these diseases it is likely mechanistic features are shared in the two contexts.

Inhibiting EZH2 has clear therapeutic potential in DMG (Mohammad et al. 2017; Piunti et al. 2017; Brien et al. 2021), as further supported by our in vivo patient-derived xenograft data in this study. Disrupting the oncogenic repression of PRC1/2 target genes reduces the differentiation blockade in DMG cells (Brien et al. 2021; Krug et al. 2021); and when combined for example with inhibitors of activated growth signalling pathways could elicit a robust therapeutic response. Tazemetostat, the current EZH2 inhibitor in approved clinical use for other cancer types has poor blood brain penetrability (Zhang et al. 2015), limiting its usefulness in the context of CNS tumours. However, strides have been made towards developing EZH2 inhibitors with optimised chemical properties to improve brain penetrability (Liang et al. 2022). Therefore, this remains a viable therapeutic avenue. However, it is noteworthy that PRC2 is a tumour suppressor in several malignancies, including childhood acute lymphoblastic leukaemia (Ernst et al. 2010; Nikoloski et al. 2010; Ntziachristos et al. 2012; Brien et al. 2016; Bracken et al. 2019). As such, systemic inhibition of PRC2 has the potential to cause secondary treatment-associated malignancies (Straining and Eighmy 2022). Our work here lays the mechanistic framework for a distinct strategy which achieves the same therapeutic endpoint i.e., activation of repressed Polycomb target genes by targeting CBX4/PCGF4 cPRC1 which we show mediate repression downstream of PRC2. Targeting specific configurations amongst the diverse cPRC1 complexes present in cells, might reduce systemic side-effects such as secondary malignancy, when compared to PRC2/EZH2 inhibition. Supporting this idea, CBX4 or PCGF4 (and indeed other cPRC1 subunits) have not been identified as tumour suppressors in somatic sequencing studies in cancer, unlike PRC2 members (Zehir et al. 2017; Bracken et al. 2019). It is noteworthy that CBX4 has a chromodomain in its N-terminus which acts as a histone lysine methylation ‘reader’ module. Reader domains have emerged as highly druggable therapeutic targets, suggesting small molecule development disrupting cPRC1 is highly feasible with some advances already made in this regard (Brien et al. 2016; Lamb et al. 2019; Suh et al. 2022).

The existence of multiple H3K27me3 readers suggests each reader contributes to distinct functional outcomes downstream of the mark. However, compelling evidence for clear functional diversity between distinct cPRC1 formations in cells is limited. Recent evidence suggests different CBXs have can have different activities related to condensate formation in vitro and potentially in cells (Plys et al. 2019; Tatavosian et al. 2019; Blackledge and Klose 2021; Brown et al. 2023). This suggests complexes defined by different CBX proteins could have different gene repressive activities. Studies mapping diverse cPRC1 formations have shown these complexes have highly overlapping chromatin binding profiles. Sequential ChIP mapping of CBX, PHC and RING paralogs even suggest distinct complexes co-bind chromatin in both space and time (Maertens et al. 2009; Pemberton et al. 2014). Indeed, we see largely overlapping binding profiles for distinct cPRC1 complexes in DMG cells. These findings imply that distinct complexes converge together at the same target sites, where their combined biochemical activities are required to mediate an appropriate level of transcriptional repression. Our discovery is a clear demonstration that within this milieu, distinct complexes do make differential functional contributions to gene repression. Our data indicate that the pattern of H3K27me3 is instrumental in correctly balancing cPRC1 repressive activities. Since H3K27me3 levels are frequently perturbed in cancer and a range of developmental diseases (Bracken et al. 2019; Deevy and Bracken 2019), it will be important to understand how altering H3K27me3 influences cPRC1 function in other disease contexts. Understanding how these complexes functionally integrate will be essential to understanding their role in normal biology, and importantly how disrupting these dynamics contributes to disease development.

## Acknowledgments

We thank members of the Brien and Bracken laboratories for helpful discussions throughout the development of this work. D.G was supported by a PhD fellowship from the Irish Research Council (IRC) Government of Ireland Postgraduate Scholarship Programme (GOIPG/2019/2084). E.L is supported by a PhD fellowship from the IRC Government of Ireland Postgraduate Scholarship Programme (GOIPG/2021/870). A.J.S is supported by a PhD fellowship from the IRC Government of Ireland Postgraduate Scholarship Programme (GOIPG/2023/4864). UCD mass spectrometry was supported by The Comprehensive Molecular Analytical Platform (CMAP) under The Science Foundation Ireland (SFI) Research Infrastructure Programme, reference 18/ RI/5702. Work in the Phillips Lab was supported by the Team Jack Foundation and the V Foundation (V2022-013). Work in the Bracken lab was supported by the Irish Cancer Society (CancersUnmetNeeds012), Worldwide Cancer Research and The Brain Tumour Charity (18-0592), an IRC Advanced Laureate Award (IRCLA/2019/21) and SFI, under the SFI Investigators program (SFI/16/IA/4562). Work in the Brien lab was supported by an SFI Starting Investigator Research Grant (18/SIRG/5573) and is currently supported by a Worldwide Cancer Research grant (21-0271) and UK Horizon Europe guarantee (ERC-2021-StG) grant.

## Author Contributions

D.G, E.L, R.E.P, A.P.B and G.L.B conceived the study. D.G performed PRC1 and PRC2 purifications, PRC2 tiling and genome-wide CRISPR screens and validations. E.L performed genome-wide CRISPR screens and validations, and all genome-wide chromatin and gene expression mapping studies. A.J.S performed PRC1 purifications and contributed to functional perturbation experiments. P.B. performed chromatin focused CRISPR screens, in vivo and in vitro PRC1 and PRC2 functional validation experiments. A.M.D, D.N. and C.M contributed to computational analyses of proteomic and genomic datasets. R.McC, F.J.S-R, Y.M.S-F and C-W.C contributed to custom CRISPR screening experiments. A.V. and J.I contributed to functional perturbation experiments. K.W, M.V and G.O performed mass spectrometry analyses. K.W, M.V and G.O performed mass spectrometry analyses. G.L.B wrote the manuscript with contributions from all authors.

## Declaration of Interests

The authors declare no competing interests.

---

**Figure S1:**
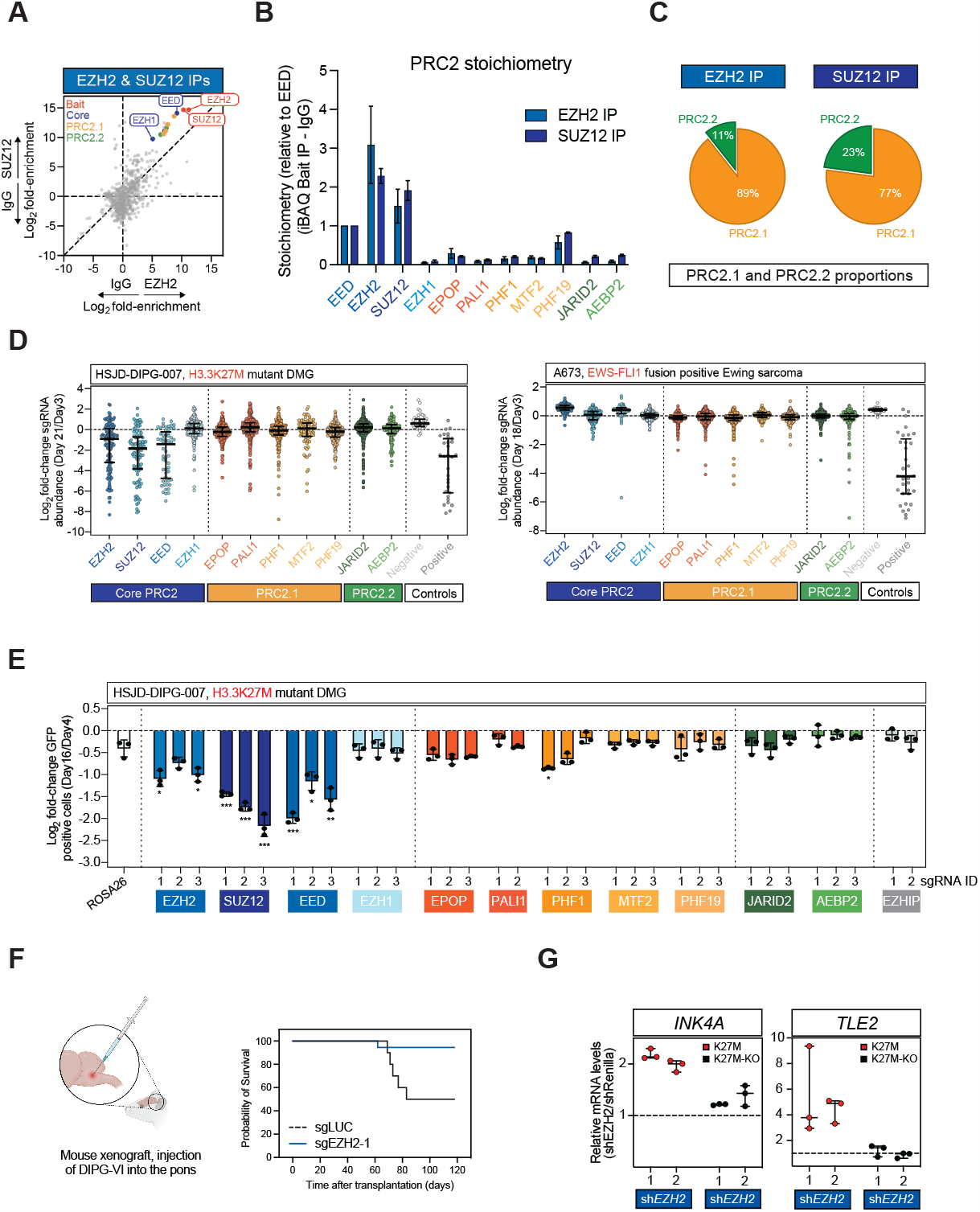
PRC2 dependency in H3K27M-mutant DMG. **A**.XY scatter plot showing log2 fold enrichment of proteins identified by mass spectrometry in endogenous EZH2 and SUZ12 immunoprecipitations. Bait (red) proteins in addition to PRC2 core (blue) and subclass defining (orange and green) are labelled and/or indicated on the plot. **B**.Bar plot showing stoichiometry of each of the indicated PRC2 member proteins in EZH2 and SUZ12 endogenous IPs. Stoichiometry of each protein is presented relative to core component EED (n = 3 pulldowns, data represents mean ± SD). **C**.Pie charts denoting the percent of total PRC2 subclass defining protein abundance (PRC2.1 and PRC2.2) identified in EZH2 and SUZ12 IP-MS. **D**.PRC2 tiling CRISPR dropout screens in HSJD-DIPG-007 cells and Ewing sarcoma A673 cells. Each dot represents the log2 fold-change (mean of n =2) in the abundance of individual sgRNAs targeting the indicated PRC2 members. The median and interquartile range of all sgRNAs targeting a given gene are indicated. **E**.Bar plots depicting log2 fold change in sgRNA abundance between day 16 and day 4 of sgRNA growth competition assays in HSJD-DIPG-007-Cas9 cells (n = 3, data represents mean ± SD). P-values calculated using Student’s t-test comparing against negative control sgRNA targeting ROSA26, * = P ≤ 0.05, ** = P ≤ 0.01 and *** = P ≤ 0.001. **F**.Survival curve for recipient mice injected with DIPGVI cells expressing control (sgLUC) or an EZH2 targeting sgRNA. **G**.Quantitative qRT-PCRs of the indicated transcripts in DIPGXIII-K27M (red) or KO (black) cells transduced with 2 independent EZH2 targeting shRNAs (n = 3, data are represented as mean ± SD). **E**. Quantitative qRT-PCRs of the indicated transcripts in DIPGXIII-K27M (red) or KO (black) cells transduced with 2 independent EZH2 targeting shRNAs (n = 3, data are represented as mean ± SD).

**Figure S2:**
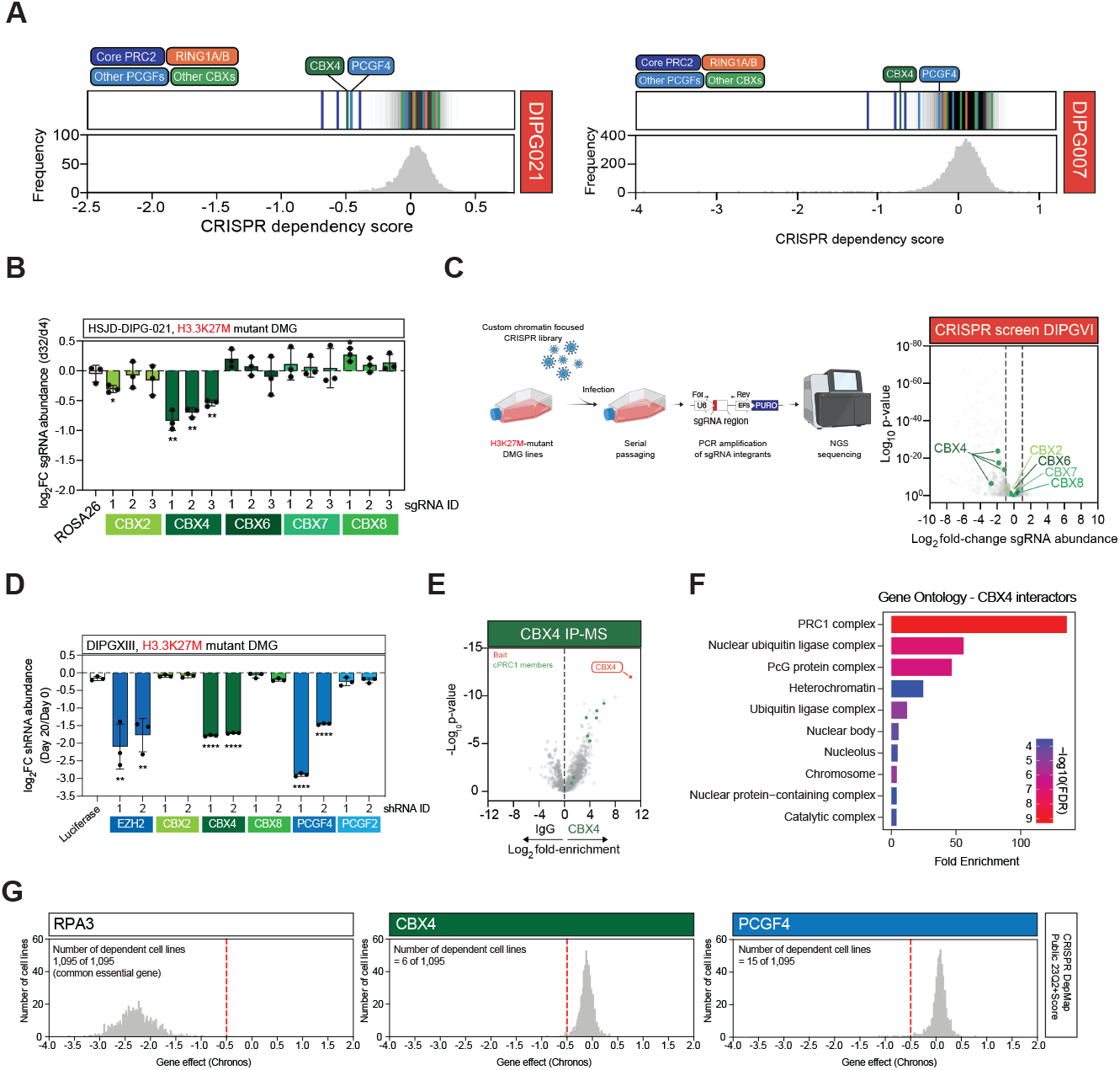
Specific functional requirement of CBX4 and PCGF4 containing cPRC1. **A**.CRISPR dependency scores calculated with the MAGeCK analysis pipeline for HSJD-DIPG-021 and HSJD-DIPG-007 cells. CBX4 and PCGF4 are indicated, with other cPRC1 and PRC2 members colour-coded on the dependency score plot. **B**.Bar plots depicting log2 fold change in sgRNA abundance between day 32 and day 4 in sgRNA growth competition assays in HSJD-DIPG-021-Cas9 cells (n = 3, data represents mean ± SD). P-values calculated using Student’s t-test comparing against negative control sgRNA targeting ROSA26, * = P ≤ 0.05 and ** = P ≤ 0.01. **C**.Schematic overview of the chromatin focused CRISPR functional screening approach in DMG cells (Left Panel); with volcano plot depicting log2 fold change in individual sgRNA abundance in DIPGVI-Cas9 cells calculated using DESeq. All sgRNAs targeting individual CBX paralogs are labelled, and colour coded. **D**.Bar plots depicting log2 fold change in shRNA abundance between day 20 and day 0 in shRNA growth competition assays in DIPGXIII-K27M cells with the data representing the mean ± SD. P-values calculated using Student’s t-test comparing against negative control shRNA targeting Luciferase, ** = P ≤ 0.01 and **** = P ≤ 0.0001. **E**.Volcano plot of endogenous CBX4 IP-MS in DIPGXIII-K27M. CBX4 and cPRC1 member proteins are colour-coded and indicated on the plot. **F**.Gene Ontology plot depicting terms enriched among proteins co-purified in endogenous CBX4 IP-MS experiments. **G**.Plots denoting the distribution of CRISPR dependency scores for the RPA3, CBX4 and PCGF4 genes across all cell lines used in the DepMap cancer dependency mapping project. The number of cell lines where each gene is denoted as a dependency (Gene Effect Score <-0.5) is indicated, relative to the total number of cell lines screened.

**Figure S3:**
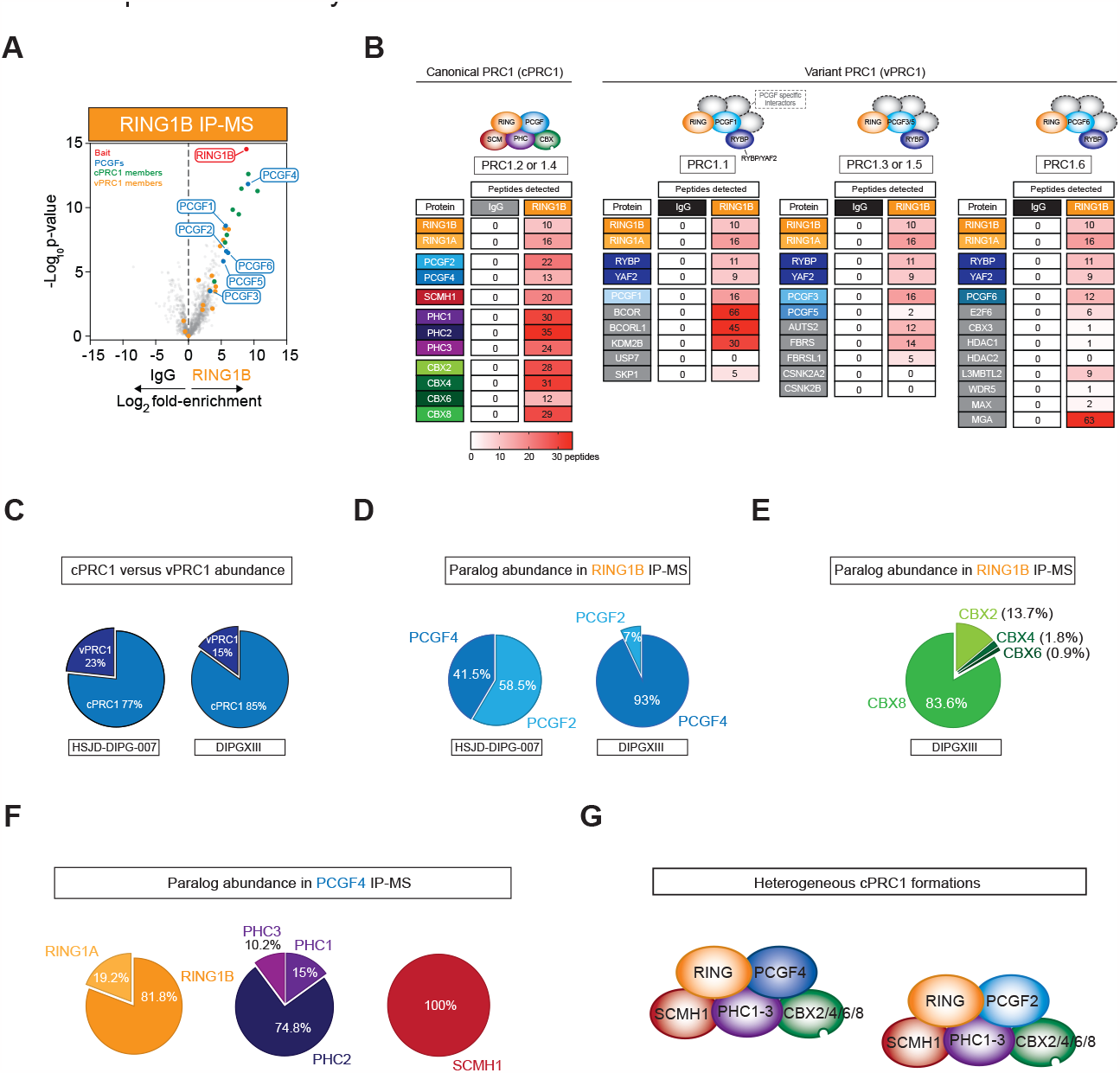
PRC1 complex biochemistry in DMG cells. **A** Volcano plot of endogenous RING1B IP-MS in DIPGXIII-K27M cells. cPRC1, vPRC1, PCGF paralogs and RING1B are colour-coded and indicated on the plot. **B**.Table depicting peptide numbers identified for each of the indicated PRC1 members in RING1B IP-MS in DIPGXIII-K27M cells, with the table separated based on PRC1 molecular subclass. **C**.Pie chart denoting the percent of total PCGF protein abundance accounted for by canonical PRC1 and variant PRC1 specific PCGF paralogs in RING1B IP-MS data in HSJD-DIPG-007 and DIPGXIII-K27M cells. **D**.Pie chart denoting the percent of total canonical PCGF protein abundance accounted for by the canonical PRC1 specific PCGF paralogs in RING1B IP-MS data in HSJD-DIPG-007 and DIPGXIII-K27M cells. **E**.Pie chart denoting the percent of total CBX protein abundance accounted for by each of the individual CBX paralogs in RING1B IP-MS in DIPGXIII-K27M cells. **F**.Pie chart denoting the percent of total protein abundance accounted for by individual proteins within each of the indicated paralog groups (RING, PHC and SCM) in PCGF4 IP-MS in DIPGXIII-K27M cells. **G**.Schematic representation of heterogenous cPRC1 make-up in DMG cells.

**Figure S4:**
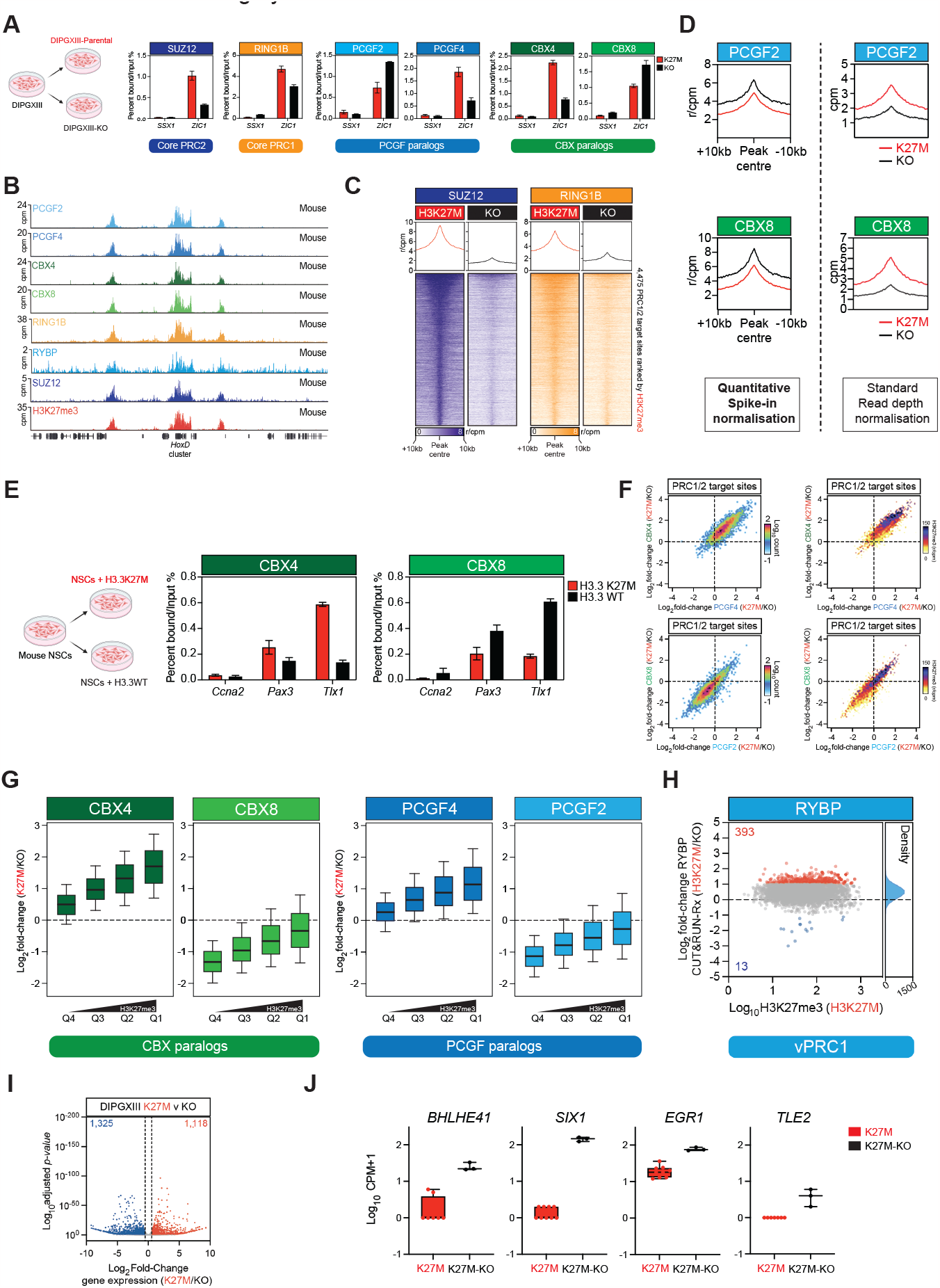
PRC1 chromatin binding dynamics. **A**.CUT&RUN-qPCRs assays for the indicated proteins in DIPGXIII-K27M (red) and DIPGXIII-KO (black) cells. The relative enrichment of each protein is demonstrated at the indicated gene promoter regions. **B**.Genomic tracks showing average PCGF2, PCGF4, CBX4, CBX8, RING1B, RYBP SUZ12 and H3K27me3 CUT&RUN signal from biological triplicate experiments at the indicated genomic locus from spike-in mouse neural stem cells. **C**.Tornado and average plots showing average enrichments of cPRC1 members in biological triplicate CUT&RUN-Rx mapping in DIPGXIII-K27M and KO cells. Indicated are the regions corresponding to the genomic window +/-10kb of PRC1/2 target sites. The scale bar denotes reference-adjusted counts per million mapped reads (r/CPM) for all experiments. **D**.Average plots denoting PCGF2 and CBX8 enrichment at PRC1/2 target sites in DIPGXIII-K27M and DIPGXIII-KO; with exogenous genome based spike-in (left panels) or read-depth based normalisation (right panels). **E**.CUT&RUN-qPCRs assays for the indicated proteins in mouse NSCs expressing H3.3K27M (red) or H3.3WT (black). The relative enrichment of each protein is demonstrated at the indicated gene promoter regions. **F**.XY scatter plot correlating changes in CBX4 and PCGF4 (top) or CBX8 and PCGF2 (bottom) binding at PRC1/2 target sites in DIPGXIII-K27M and KO cells (left); overlay of H3K27me3 signal in DIPGXIII-K27M at PRC1/2 targets as per left panels. **G**.Boxplots showing log2 fold change of CBX4, CBX8, PCGF4 and PCGF2 chromatin binding between DIPGXIII-K27M and KO cells at PRC1/2 target regions; sites are separated into quartiles based on H3K27me3 enrichment in DIPGXIII-K27M cells. **H**.MA-plots showing fold-change in binding for RYBP at PRC1/2 target sites in DIPGXIII-K27M and KO cells. Sites changing >2-fold are shown in red (increasing) or blue (decreasing) with the number of sites in these categories also indicated. The densities of chromatin binding changes are shown on the right of the plot. **I**.Volcano plot depicting gene expression changes between DIPGXIII-K27M and KO cells. Number of differentially expressed genes are indicated on the plot. **J**.Boxplots presenting expression values for the indicated PRC1/2 target genes in DIPGXIII-K27M and KO cells (n = 3 biological replicates).

**Figure S5:**
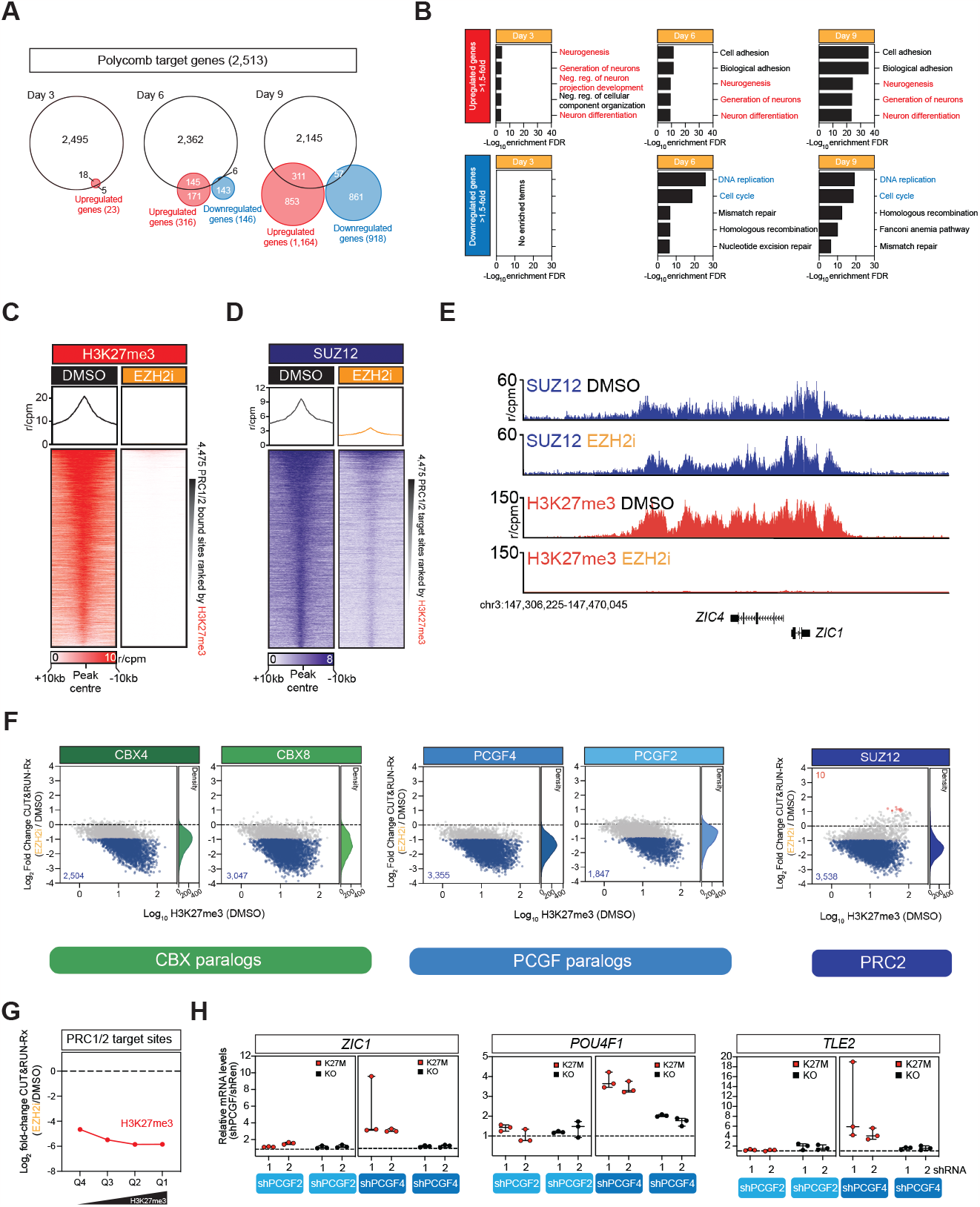
Specific transcriptional role for CBX4 and PCGF4 containing complexes. **A**.Venn diagrams depicting the total number of PRC1/2 target genes and their overlap with genes differentially expressed following 3, 6 and 9 days of EZH2 inhibition. **B**.Gene Ontology analyses of differentially expressed genes following 3, 6 and 9 days of EZH2 inhibition. Depicted are the top 5 categories enriched among up or down regulated genes at each time point. **C, D**. Tornado and average plots showing average enrichments of H3K27me3 (C) and SUZ12 (D) in biological triplicate CUT&RUN-Rx mapping in DIPGXIII-K27M cells treated with EZH2 inhibitor for 6 days. Indicated are the regions corresponding to the genomic window +/-10kb of PRC1/2 target sites. The scale bar denotes reference-adjusted counts per million mapped reads (r/CPM) for all experiments. **E**.Genomic tracks showing average quantitative SUZ12 and H3K27me3 CUT&RUN-Rx signal from biological triplicate experiments at the indicated genomic locus in control (DMSO) treated DIPGXIII-K27M or cells treated with EZH2 inhibitor for 6 days. **F**.MA-plots showing fold-change in binding for CBX4, CBX8, PCGF4, PCGF2 and SUZ12 at PRC1/2 target sites in DIPGXIII-K27M treated with EZH2 inhibitor for 6 days. Sites changing >2-fold are shown in red (increasing) or blue (decreasing) with the number of sites in these categories also indicated. The densities of chromatin binding changes are shown on the right of each plot. **G**.Dynamics of H3K27me3 changes at PRC1/2 target gene quartiles in DIPGXIII-K27M cells treated with EZH2 inhibitor for 6 days. Shown are the mean changes relative to control (DMSO) treated cells. **H**.Quantitative qRT-PCRs of the indicated transcripts in DIPGXIII-K27M (red) or KO (black) cells transduced with 2 independent shRNAs targeting PCGF2 or PCGF4 (n = 3, data represents mean ± SD).

## EXPERIMENTAL MODEL AND SUBJECT DETAILS

### Cell Culture and Lentiviral Production

All cell cultures were maintained in a humidified, normoxic incubator at 37°C, 5% CO2. Patient-derived HSJD-DIPG-007, HSJD-DIPG-021, DIPGXIII-K27M and DIPGXIII-KO cells were grown as monolayers. SU-DIPGVI, SU-DIPG-XIII, SU-DIPG50, GBM667 were grown as neurospheres. Monolayer cultures were grown in Neural Stem Cell (NSC) media composed of DMEM/F-12 media (Sigma) supplemented with 8mM Glucose (Sigma), 100mM MEM Non-Essential Amino Acids (Gibco), 100U/ml Penicillin/100mg/ml Streptomycin (Gibco), 0.12g/l BSA (Gibco), 100mM β-mercaptoethanol (Gibco), 1% B27 (Gibco), 0.5% N2 (Gibco), 10ng/ml mouse EGF (Peprotech), 10ng/ml human FGF (Peprotech) and 2-4μg/ml Laminin (Cultrex), or in Tumour Stem Media (TSM) media composed of 50:50 DMEM/F12 and Neurobasal-A media supplemented with 1% HEPES 1 M (Gibco), 100mM MEM Non-Essential Amino Acids (Gibco), 1 mM Sodium Pyruvate (Gibco), 1% GlutaMAX (Gibco), 2% B27 without vitamin A (Gibco), 20ng/ml human EGF (Peprotech), 20ng/ml human FGF (Peprotech), 10ng/ml human PDGF-AA (Peprotech), 10ng/ml human PDGF-BB (Peprotech), 5 IU/ml Heparin (Sigma) and 2-4μg/ml Laminin (Cultrex). For neurosphere culture Laminin was omitted from NSC media supplemented with 20ng/ml EGF, 20ng/ml FGF, 10ng/ml PDGF-AA and 10ng/ml PDGF-BB. Cells were passaged every 3-5 days using Accutase.

Lentiviral packaging HEK293T and A673 (Ewing sarcoma) cells were cultured in DMEM media (Sigma) supplemented with 10% foetal bovine serum and 100 U/ml Penicillin/100 μg/ml Streptomycin (Gibco). Lentiviral particles were generated by cotransfection of HEK293T cells with a lentiviral expression vector (sgRNA/shRNA/cDNA) or pooled sgRNA library with a viral packaging (PAX2) and envelope (VSV-G) vectors using PEI or X-tremegene (Roche) in accordance with standard protocols. Viral supernatants were collected between 24-72hrs posttransfection and either used directly for infection of target cells after filtering through a 0.45mm syringe filter and addition of 8.5 mg/ml Polybrene; or concentrated by ultracentrifugation (20,000 rpm, 2hrs) before being resuspended in 0.5-1ml of neural stem cell culture media and used for infection of target cells.

### Endogenous immunoprecipitations

Harvested cells were washed twice with cold PBS and washed pellets resuspended with Buffer A (25mM HEPES pH 7.6, 5mM MgCl2, 25mM KCl, 0.05mM EDTA, 10% (v/v) glycerol, 0.1% (v/v) NP40, 1mM DTT and 1mM PMSF) and kept on ice for 10 minutes. Samples were centrifuged at 1,500rpm for 10 minutes at 4°C and the supernatant was discarded. To lyse nuclei, pellets were resuspended in Buffer C (10mM HEPES pH 7.6, 3mM MgCl2, 100mM KCl, 0.5mM EDTA, 10% (v/v) glycerol, 1mM DTT and 1mM PMSF). Next, 10% (v/v) 3M (NH4)2SO4 prepared in Buffer C was added to the samples, which were rotated at 4°C for 20 minutes. Samples were ultracentrifuged at 350,000 rcf for 15 minutes at 4°C in a Beckman Coulter Optima L-100XP using a SW 55Ti rotor. The supernatant was collected and 300mg of (NH4)2SO4 was added for each 1mL of sample, to precipitate nuclear extracts. Samples were vortexed and kept on ice for 15 minutes before being ultracentrifuged using the same parameters as above. Pellets were resuspended in IP buffer (300mM NaCl, 50mM Tris-HCl pH 7.5, 1mM EDTA, 1% (v/v) Triton-X-100, 1mM DTT and 1mM PMSF) and Bradford assays were performed to determine protein concentration. Benzonase Nuclease (Sigma) and 1μg of the respective antibody were added to each IP and samples were rotated at 4°C overnight. The following day, Protein A Sepharose 4B, Fast Flow beads (Sigma) (for PRC2 experiments) or Protein G Dynabeads (Thermo Fisher) (for PRC1 experiments) were added to each sample, followed by rotation at 4°C for 2.5 hours. Beads were washed 5x with IP buffer, before being processed for immunoblotting or mass spectrometry sample preparation.

### PRC2 mass spectrometry

Beads were resuspended in elution buffer (2M Urea, 100 mM Tris pH 8, 10 mM DTT) and incubated 20 min on a shaker (1300 rpm) at RT. After incubation, iodoacetamide was added to a final concentration of 50 mM, followed by 10 min shaking in the dark at RT. Partial digestion and elution from the beads was initiated by adding 0.25 mg Trypsin (Promega; V5113) for 2 hr. The supernatant containing the IP samples was collected and the beads were resuspended in 50 μL elution buffer followed by a 5 min incubation shaking at RT. Both supernatants were combined, and 0.1 mg Trypsin was added followed by overnight incubation at RT. The digestion was stopped by adding TFA (final concentration 0.5%). The resulting digested samples were desalted and purified using StageTips(Rappsilber et al. 2007). The peptides were eluted from StageTips with buffer B (80% acetonitrile, 0.1% formic acid), concentrated to 5 μL by SpeedVac centrifugation at room temperature, and filled up to 12 μL using buffer A (0.1% formic acid). Pull-down samples were measured using a gradient from 9%–32% Buffer B for 114 min followed by washes at 50% then 95% Buffer B resulting in total 140 min data collection time. Mass spectra were recorded on an LTQ-Orbitrap Fusion Tribrid mass spectrometer (Thermo Fisher Scientific). Scans were collected in data-dependent top speed mode with dynamic exclusion set at 60 s.

### PRC1 mass spectrometry

In-solution tryptic digestions were performed as described previously(Wisniewski et al. 2009). Samples were run on a Bruker timsTof Pro mass spectrometer connected to an Evosep One liquid chromatography system. Tryptic peptides were resuspended in 0.1% formic acid and each sample was loaded on to an Evosep tip and separated on a Evosep EV1137 Performance Column – 15 cm x 150 um, 1.5 um. The Evosep tips were placed in position on the Evosep One, in a 96-tip box. The autosampler is configured to pick up each tip, elute and separate the peptides using a set chromatography method (15 samples a day)(Bache et al. 2018). The mass spectrometer was operated in positive ion mode with a capillary voltage of 1650 V, dry gas flow of 3 l/min and a dry temperature of 180 °C. All data was acquired with the instrument operating in trapped ion mobility spectrometry (TIMS) mode. Trapped ions were selected for ms/ms using parallel accumulation serial fragmentation (PASEF). A scan range of (100-1700 m/z) was performed at a rate of 5 PASEF MS/MS frames to 1 MS scan with a cycle time of 1.03s(Meier et al. 2018).

### Design and synthesis of human PRC2 sgRNA library

For the PRC2 complex member gene tiling CRISPR library, a total of 3,599 sgRNA sequences targeting the coding exons of the PRC2 complex genes (*AEBP2, EED, EPOP, EZH1, EZH2, JARID2, PALI1, MTF2, PHF1, PHF19* and *SUZ12*) were designed using the Genetic Perturbation Platform (Broad Institute)(Doench et al. 2016). Briefly, guide RNA oligos were synthesized by microarray (CustomArray) and cloned into the ipUSEPR lentiviral sgRNA vector (hU6-driven sgRNA co-expressed with EF-1a-driven red fluorescent protein [RFP] and puromycin-resistance gene) using the BsmBI (NEB) restriction sites. For a complete list of PRC2 scanning library sgRNA sequences see Table S1. Individual sgRNA selected for validation experiments are listed in Table S4. CRISPR lentivirus was produced in HEK293 cells as described above.

### Tiled PRC2 CRISPR screen

Cas9-expressing HSJD-DIPG-007, HSJD-DIPG-021 and A673 cells were transduced with the PRC2 sgRNA lentiviral library at a multiplicity of infection (MOI) between 0.1-0.3 at 1000x library coverage. Approximately 24 hours following transduction, puromycin was added to cell cultures to select for sgRNA expressing cells. Cell pellets were harvested for genomic DNA extraction at Day 3 and Day 18-21 post-transduction. Genomic DNA was extracted using the Qiagen QIAmp DNA Blood Mini Kit, following the manufacturer’s protocol. PCR using ipUSEPR specific primers (DCF01 5’-CTTGTGGAAAGGACGAAACACCG-3’ and DCR03 5’-CCTAGGAACAGCGGTTTAAAAAAGC-3’) was performed: 50μL DreamTaq Hot Start Green PCR Master Mix (Thermo Scientific), 0.5μL P5 primer mix (100μM), 0.5μL P7 index primer (100μM) using the following cycling conditions: 95°C for 60s, 28 cycles of 95°C for 30s, 53°C for 30s, 72°C for 30s; 72°C for 10min. Multiple PCRs on 2.5-5 μg input gDNA were performed for each gDNA pool. PCR products were subject to agarose gel electrophoresis, gel purified, and isopropanol precipitated. Individually indexed libraries were pooled and sequenced using an Illumina NextSeq500. Primers used to prepare sequencing libraries are shown in Table S2.

### Genome-wide Brunello CRISPR screen

Cas9-expressing HSJD-DIPG-007 and HSJD-DIPG-021 cells were transduced with the Brunello lentiviral library at a target MOI ∼0.3 at 350-500x library coverage. Approximately 24 hours following transduction, puromycin was added to cell cultures to select for sgRNA expressing cells. Cell pellets were harvested for genomic DNA extraction at Day 21 or Day 28 post-transduction. Genomic DNA was extracted using the NucleoSpin Blood L (Macherey-Nagel), following the manufacturer’s protocol. Prior to PCR, the OneStep PCR Inhibitor Removal Kit (Zymo Research) was used, following the manufacturer’s protocol. PCR using lentiGuide-Puro specific primers (from Broad GPP, Supplementary Table X) and PCR purification was performed as per tiled pooled PRC2 screen above. Individually indexed libraries were pooled and sequenced using an Illumina NextSeq500.

### Chromatin focused CRISPR screen

Cas9-expressing SU-DIPG-VI cells were transduced with a custom sgRNA library containing sgRNAs targeting ∼550 genes cloned into the ipUSEPR vector backbone as previously described(Phillips et al. 2019; Soto-Feliciano et al. 2023). An MOI of ∼0.2 at 1000x library coverage was used. 5 days following transduction, puromycin was added to cell cultures to select for sgRNA expressing cells. Cell pellets were harvested for genomic DNA extraction at approximately 15 cumulative population doublings post-transduction. Genomic DNA was extracted using the Plasmid Plus Maxi Kit (Qiagen). We used a modified two-step PCR version of the protocol published by Doench and colleagues(Doench et al. 2016). Briefly, we performed an initial “enrichment” PCR, whereby the integrated sgRNA cassettes were amplified from gDNA, followed by a second PCR to append Illumina sequencing adapters on the 5′- and 3′-ends of the amplicon, as well as a random nucleotide stagger and unique demultiplexing barcode on the 5′-end. Each “PCR1” reaction contains 25 μL of Q5 High-Fidelity 2X Master Mix (NEB), 2.5 μL of Nuc PCR#1 Fwd Primer (10 μm), 2.5 μL of Nuc PCR#1 Rev Primer (10 μmol/L), and 5 μg of gDNA in 20 μL of water (for a total volume of 50 μL per reaction). The number of PCR1 reactions is scaled-accordingly; therefore, we performed six PCR1 reactions per technical replicate, per time point (T0 or TF). PCR1 amplicons were purified using the QIAquick PCR Purification Kit (Qiagen) and used as a template for “PCR2” reactions. Each PCR2 reaction contains 25 μL of Q5 High-Fidelity 2X Master Mix (NEB), 2.5 μL of a unique Nuc PCR#2 Fwd Primer (10 μmol/L), 2.5 μL of Nuc PCR#2 Rev Primer (10 μmol/L), and 300 ng of PCR1 product in 20 μL of water (for a total volume of 50 μL per reaction). We performed two PCR2 reactions per PCR1 product. Library amplicons were size-selected on a 2.5% agarose gel, purified using the QIAquick Gel Extraction Kit (Qiagen), and sequenced on an Illumina NextSeq 500 instrument (100-nt, single-end reads).

### Negative selection assays

Cas9-expressing DMG cell lines were transduced with sgRNA-GFP or RFP expressing lentivirus at a low MOI and passaged without selection. The percentage of GFP or RFP-positive cells was measured at each passage using the Guava easyCyte Flow Cytometer (Cytek) or LSRII (BD). The relative proportion of GFP/RFP positive cells was monitored throughout this serial cell culture. For shRNA negative selection experiments DMG cells were transduced with shRNA-GFP expressing lentivirus and selected with puromycin. Selected cultures were mixed 50:50 with non-transduced (GFP negative) cells and the percentage of GFP positive cells was monitored at each passage using the Guava easyCyte Flow Cytometer (Cytek). Sequences of sgRNAs and shRNAs used in these experiments can be found in Table S4 and S5.

### CUT&RUN-Rx

Cells were harvested using Accutase (STEMCELL Technologies) and counted. Following counting mouse NSCs were spiked-in to human DIPGXIII cell suspensions (1:10 cell number ratio). Cells were fixed in culture media containing 0.1% formaldehyde at room temperature for 1 min. Formaldehyde was quenched with glycine at 0.125 M, followed by a 5-min incubation at room temperature. Fixed cells were washed with PBS. Fixed cells were incubated on ice for 10 min in nuclear extraction buffer (20mM HEPES pH 7.5, 10mM KCL, 0.1% Triton X-100, 20% Glycerol,1X protease inhibitor cocktail and 0.5mM spermidine). Extracted nuclei were collected by centrifugation (4ºC, 600g) and resuspended in cold nuclei extraction buffer (100μL per sample). CUT&RUN was performed following Epicypher CUT&RUN Protocol v1.5.1. CUT&RUN DNA was purified using the Monarch PCR & DNA Cleanup Kit (New England Biolabs) in accordance with the manufacturer’s instructions. Primer sequences used for quantitative PCR analysis can be found in Table S6.

### CUT&RUN-Rx Library Preparation

Purified DNA was quantified using a Qubit fluorimeter (Invitrogen). 1-10 ng of DNA was used to generate CUT&RUN-Rx libraries with the NEBNext Ultra II DNA Library Prep Kit for Illumina (New England Biolabs) as per the manufacturer’s instructions. Library DNA was quantified using the Qubit, and size distributions were ascertained on a TapeStation (Agilent) using the D1000 ScreenTape assay reagents (Agilent; 5067-5583). This information was used to calculate pooling ratios for multiplex library sequencing. Pooled libraries were diluted and processed for 38-bp paired-end sequencing on an Illumina NextSeq500 instrument using the NextSeq 500 High Output v2 kit (Illumina; FC-404-2005) in accordance with the manufacturer’s instructions.

### Immunoblotting

Whole cell protein samples were prepared in RIPA buffer (25 mM Tris-HCl. pH7.6, 150 mM NaCl, 1% NP-40, 1% Sodium Deoxycholate, 0.1% SDS) containing a 1X protease inhibitor cocktail. Protein lysates were separated on pre-cast Bolt 4–12% Bis-Tris Plus Gels (Invitrogen, NW04127BOX) and transferred to nitrocellulose membranes. Membranes were probed using the relevant primary and secondary antibodies and relative protein levels were determined using the Odyssey CLx or Fc Imager (LI-COR).

### Quant-seq sample prep and sequencing

Total RNA was isolated from untreated DIPGXIII-K27M/KO cells, or DIPGXIII-K27M following 3, 6 or 9 days of Tazemetostat (2μM) treatment. The quality of extracted RNA was confirmed using the TapeStation (Agilent) with the RNA ScreenTape assay reagents (Agilent; 5067-5576). Total RNA (500 ng) was used as input material from each sample for library preparation. Libraries were generated using the QuantSeq 3′ mRNA-Seq Library Prep Kit FWD for Illumina (Lexogen; 015.24) in accordance with the manufacturer’s instructions. Library DNA was quantified using the Qubit, and size distributions were ascertained on a TapeStation (Agilent) using the D1000 ScreenTape assay reagents (Agilent; 5067-5583). This information was used to calculate pooling ratios for multiplex library sequencing. Pooled libraries were diluted and processed for 75-bp single-end sequencing on an Illumina NextSeq instrument using the NextSeq 500 High Output v2 kit (75 cycles) (Illumina; FC-404-2005) in accordance with the manufacturer’s instructions.

### Quantitative RT-PCR

Total RNA was extracted from cells using the RNeasy kit (Qiagen) according to the manufacturer’s protocol. RNA was used to generate cDNA by reverse transcriptase PCR using the High-Capacity cDNA Reverse Transcription Kit (Applied Biosystems). Relative mRNA expression levels were determined using SYBR Green I detection chemistry (Applied Biosystems) on the QuantStudio instrument. Primer sequences used in quantitative experiments can be found in Table S6.

### Xenograft Experiments

All mouse *in vivo* experiments were approved by the Institutional Animal Care and Use Committee (IACUC) at the University of Pennsylvania and the Memorial Sloan Kettering Cancer Center. Intracranial xenografts were established by stereotactic injection (Stoelting) of sgRNA/Cas9 expressing SU-DIPG-VI cells into the brainstem of NSG recipient mice at the age of 6-8 weeks as previously described(Monje et al. 2011). Tumor engraftment/burden was determined via either MRI or *in vivo* luminescence imaging (IVIS Spectrum, PerkinElmer), and total flux was calculated by included software (Living Image, Perkin Elmer) as the radiance through standard circular ROIs centered on the animal’s head. Experiment endpoint was triggered by morbidity criteria for euthanasia. The investigators were blinded to animals group at time of outcome assessment. Kaplan-Meier survival curves of mice and the Log-rank test were preformed using the Prism 6 software.

## QUANTIFICATION AND STATISTICAL ANALYSIS

### CRISPR screen data analysis

For the PRC2 tiling screen, sgRNA read counts were generated using PinAPL-py(Spahn et al. 2017). For scatter plots presenting log2 fold-change of sgRNA abundance, any sgRNAs with Off-target Tier I and II Bin 1 scores > 0 were removed and for a given cell line, any sgRNAs with less than 50 reads in any Day 3 replicate were also removed. Log2 fold-changes for each sgRNA were calculated by comparing the number of reads mapping to individual sgRNAs at day 3 and day 18/21 normalised to library sequencing depth. Plots were generated using Prism. For the genome-wide CRISPR screens, the MAGeCK algorithm was used to calculate CRISPR dependency beta scores comparing sgRNA abundance in the original plasmid pool with gDNA extracted at the end of a screen, using the count and mle functions(Wang et al. 2019). In the genome-wide CRISPR dependency score plots, the frequency of CRISPR dependency scores was calculated by rounding beta scores for each gene to two decimal places and counting the number of times each of the rounded scores was present. Common essential genes, as defined by depmap.org (Achilles, 19Q3), were removed and plots were generated using Prism. See Table S3 for a list of beta scores calculated from genome-wide screening experiments.

### Mass spectrometry data analysis

#### PRC2 IP-MS analysis

Mass spectrometry analysis Thermo RAW files were searched against the Homo sapiens subset of the Uniprot Swissprot database (reviewed) using the search engine Maxquant. Additional parameters that were enabled were match-between-runs, label-free quantification (LFQ) and IBAQ. After filtering for proteins that were present at least in all replicates of one condition, LFQ values were log2 transformed and missing values were imputed in Perseus using default parameters (width = 0.3, shift = 1.8). Statistical outliers for the pull-downs were determined using a two-tailed t test. Multiple testing correction was performed using a permutation-based false discovery rate (FDR) method in Perseus. For these analyses PALI1/2 were manually added to the UniProt database.

#### PRC1 IP-MS analysis

All mass spectrometry data was processed using MaxQuant software using the human UniProt database. The raw data was searched against the Homo sapiens subset of the Uniprot Swissprot database (reviewed) using the search engine Maxquant (Version 2.0.3.0) using specific parameters for trapped ion mobility spectra data dependent acquisition (TIMS DDA). Each peptide used for protein identification met specific Maxquant parameters, i.e., only peptide scores that corresponded to a false discovery rate (FDR) of 0.01 were accepted from the Maxquant database search. The normalised protein intensity of each identified protein was used for label free quantitation (LFQ)(Cox et al. 2014).

The generated ‘proteingroups.txt’ files were filtered for contaminants in R. The DEP package in R was used to make volcano plots. iBAQ intensities were used to determine the stoichiometry of the identified PRC1/PRC2 members as previously described(Smits et al. 2013).

#### CUT&RUN-Rx data analysis

Firstly, the prefix mm10\_ was added to the chromosome names of the mm10 genome build. Next, a human/mouse hybrid genome was generated by appending the human hg38 genome build to the mm10 genome build prior to indexing with bowtie2. Bowtie2 was used to align reads to the hybrid genome in the --very-sensitive mode. SAMtools was employed to retain reads with a mapping quality >2, separate the human and mouse reads, remove duplicate reads and convert SAM files to BAM files. The spikein normalisation factor for each target was calculated using the formula for normalised reference-adjusted reads per million (RRPM) (1 per million spike-in reads). The bamCoverage module from the deepTools suite was used to generate RRPM normalised bigwig files for genome browser visualisation with the following parameters where SCALEFACTOR is equal to the respective RRPM normalization factor. Peaks were called using macs2, BroadPeak files were converted to bed format using standard bash commands. The bedtools intersect module was used to exclude regions that overlapped with a custom hg38 blacklist, and to overlap peaks to generate consensus peak sets from replicate experiments. Peaks were annotated with HOMER.

### Quant-seq data analysis

The BBDuk module from the BBTools package was used to perform adapter trimming and quality filtering as recommended by the manufacturer. The STAR aligner was used to align filtered reads to the hg38 human genome build. The htseq-count tool was used to calculate the number of reads that align to each gene. Differentially expressed genes were detected using a >1.5 fold change and a <0.05 BenjaminiHochberg adjusted P value using the DESeq2 R package.

## Notes

### Competing Interest Statement

The authors have declared no competing interest.

## REFERENCES

Bache N, Geyer PE, Bekker-Jensen DB, Hoerning O, Falkenby L, Treit PV, Doll S, Paron I, Muller JB, Meier F et al. 2018. A Novel LC System Embeds Analytes in Pre-formed Gradients for Rapid, Ultra-robust Proteomics. Mol Cell Proteomics 17: 2284–2296.

Balakrishnan I, Danis E, Pierce A, Madhavan K, Wang D, Dahl N, Sanford B, Birks DK, Davidson N, Metselaar DS et al. 2020. Senescence Induced by BMI1 Inhibition Is a Therapeutic Vulnerability in H3K27M-Mutant DIPG. Cell Rep 33: 108286.

Bender S, Tang Y, Lindroth AM, Hovestadt V, Jones DT, Kool M, Zapatka M, Northcott PA, Sturm D, Wang W et al. 2013. Reduced H3K27me3 and DNA hypomethylation are major drivers of gene expression in K27M mutant pediatric high-grade gliomas. Cancer Cell 24: 660–672.

Bernstein E, Duncan EM, Masui O, Gil J, Heard E, Allis CD. 2006. Mouse polycomb proteins bind differentially to methylated histone H3 and RNA and are enriched in facultative heterochromatin. Mol Cell Biol 26: 2560–2569.

Blackledge NP, Klose RJ. 2021. The molecular principles of gene regulation by Polycomb repressive complexes. Nat Rev Mol Cell Biol 22: 815–833.

Bracken AP, Brien GL, Verrijzer CP. 2019. Dangerous liaisons: interplay between SWI/SNF, NuRD, and Polycomb in chromatin regulation and cancer. Genes Dev 33: 936–959.

Bracken AP, Dietrich N, Pasini D, Hansen KH, Helin K. 2006. Genome-wide mapping of Polycomb target genes unravels their roles in cell fate transitions. Genes Dev 20: 1123–1136.

Brien GL, Bressan RB, Monger C, Gannon D, Lagan E, Doherty AM, Healy E, Neikes H, Fitzpatrick DJ, Deevy O et al. 2021. Simultaneous disruption of PRC2 and enhancer function underlies histone H3.3-K27M oncogenic activity in human hindbrain neural stem cells. Nat Genet 53: 1221–1232.

Brien GL, Valerio DG, Armstrong SA. 2016. Exploiting the Epige-nome to Control Cancer-Promoting Gene-Expression Programs. Cancer Cell 29: 464–476.

Brown K, Chew PY, Ingersoll S, Espinosa JR, Aguirre A, Espinoza A, Wen J, Astatike K, Kutateladze TG, Collepardo-Guevara R et al. 2023. Principles of assembly and regulation of condensates of Polycomb repressive complex 1 through phase separation. Cell Rep 42: 113136.

Castel D, Kergrohen T, Tauziede-Espariat A, Mackay A, Ghermaoui S, Lechapt E, Pfister SM, Kramm CM, Boddaert N, Blauwblomme T et al. 2020. Histone H3 wild-type DIPG/DMG overex-pressing EZHIP extend the spectrum diffuse midline gliomas with PRC2 inhibition beyond H3-K27M mutation. Acta Neuropathol 139: 1109–1113.

Conway E, Healy E, Bracken AP. 2015. PRC2 mediated H3K27 methylations in cellular identity and cancer. Curr Opin Cell Biol 37: 42–48.

Cox J, Hein MY, Luber CA, Paron I, Nagaraj N, Mann M. 2014. Accurate proteome-wide label-free quantification by delayed normalization and maximal peptide ratio extraction, termed Max-LFQ. Mol Cell Proteomics 13: 2513–2526.

Deevy O, Bracken AP. 2019. PRC2 functions in development and congenital disorders. Development 146.

Doench JG, Fusi N, Sullender M, Hegde M, Vaimberg EW, Donovan KF, Smith I, Tothova Z, Wilen C, Orchard R et al. 2016. Optimized sgRNA design to maximize activity and minimize off-target effects of CRISPR-Cas9. Nat Biotechnol 34: 184–191.

Ernst T, Chase AJ, Score J, Hidalgo-Curtis CE, Bryant C, Jones AV, Waghorn K, Zoi K, Ross FM, Reiter A et al. 2010. Inactivating mutations of the histone methyltransferase gene EZH2 in mye-loid disorders. Nat Genet 42: 722–726.

Fellmann C, Hoffmann T, Sridhar V, Hopfgartner B, Muhar M, Roth M, Lai DY, Barbosa IA, Kwon JS, Guan Y et al. 2013. An optimized microRNA backbone for effective single-copy RNAi. Cell Rep 5: 1704–1713.

Feng J, Liu T, Qin B, Zhang Y, Liu XS. 2012. Identifying ChIP-seq enrichment using MACS. Nat Protoc 7: 1728–1740.

Filbin MG, Tirosh I, Hovestadt V, Shaw ML, Escalante LE, Mathewson ND, Neftel C, Frank N, Pelton K, Hebert CM et al. 2018. Developmental and oncogenic programs in H3K27M gli-omas dissected by single-cell RNA-seq. Science 360: 331–335.

Fontebasso AM, Papillon-Cavanagh S, Schwartzentruber J, Nikbakht H, Gerges N, Fiset PO, Bechet D, Faury D, De Jay N, Ramkissoon LA et al. 2014. Recurrent somatic mutations in ACVR1 in pediatric midline high-grade astrocytoma. Nat Genet 46: 462–466.

Francis NJ, Kingston RE, Woodcock CL. 2004. Chromatin compaction by a polycomb group protein complex. Science 306: 1574–1577.

Gao Z, Zhang J, Bonasio R, Strino F, Sawai A, Parisi F, Kluger Y, Reinberg D. 2012. PCGF homologs, CBX proteins, and RYBP define functionally distinct PRC1 family complexes. Mol Cell 45: 344–356.

Glancy E, Ciferri C, Bracken AP. 2021. Structural basis for PRC2 engagement with chromatin. Curr Opin Struct Biol 67: 135–144.

Glancy E, Wang C, Tuck E, Healy E, Amato S, Neikes HK, Mariani A, Mucha M, Vermeulen M, Pasini D et al. 2023. PRC2.1- and PRC2.2-specific accessory proteins drive recruitment of different forms of canonical PRC1. Mol Cell 83: 1393–1411 e1397.

Grobner SN, Worst BC, Weischenfeldt J, Buchhalter I, Kleinheinz K, Rudneva VA, Johann PD, Balasubramanian GP, Segura-Wang M, Brabetz S et al. 2018. The landscape of genomic alterations across childhood cancers. Nature 555: 321–327.

Hanahan D. 2022. Hallmarks of Cancer: New Dimensions. Cancer Discov 12: 31–46.

Harutyunyan AS, Krug B, Chen H, Papillon-Cavanagh S, Zeinieh M, De Jay N, Deshmukh S, Chen CCL, Belle J, Mikael LG et al. 2019. H3K27M induces defective chromatin spread of PRC2-me-diated repressive H3K27me2/me3 and is essential for glioma tu-morigenesis. Nat Commun 10: 1262.

Hauri S, Comoglio F, Seimiya M, Gerstung M, Glatter T, Hansen K, Aebersold R, Paro R, Gstaiger M, Beisel C. 2016. A High-Density Map for Navigating the Human Polycomb Complexome. Cell Rep 17: 583–595.

Jain SU, Do TJ, Lund PJ, Rashoff AQ, Diehl KL, Cieslik M, Bajic A, Juretic N, Deshmukh S, Venneti S et al. 2019. PFA ependymo-ma-associated protein EZHIP inhibits PRC2 activity through a H3 K27M-like mechanism. Nat Commun 10: 2146.

Jain SU, Rashoff AQ, Krabbenhoft SD, Hoelper D, Do TJ, Gibson TJ, Lundgren SM, Bondra ER, Deshmukh S, Harutyunyan AS et al. 2020. H3 K27M and EZHIP Impede H3K27-Methylation Spreading by Inhibiting Allosterically Stimulated PRC2. Mol Cell 80: 726–735 e727.

Jessa S, Blanchet-Cohen A, Krug B, Vladoiu M, Coutelier M, Faury D, Poreau B, De Jay N, Hebert S, Monlong J et al. 2019. Stalled developmental programs at the root of pediatric brain tumors. Nat Genet 51: 1702–1713.

Jones C, Baker SJ. 2014. Unique genetic and epigenetic mechanisms driving paediatric diffuse high-grade glioma. Nat Rev Cancer 14.

Jovanovich N, Habib A, Head J, Hameed F, Agnihotri S, Zinn PO. 2023. Pediatric diffuse midline glioma: Understanding the mechanisms and assessing the next generation of personalized therapeutics. Neurooncol Adv 5: vdad040.

Kagey MH, Melhuish TA, Wotton D. 2003. The polycomb protein Pc2 is a SUMO E3. Cell 113: 127–137.

Khuong-Quang DA, Buczkowicz P, Rakopoulos P, Liu XY, Fontebasso AM, Bouffet E, Bartels U, Albrecht S, Schwartzentruber J, Letourneau L et al. 2012. K27M mutation in histone H3.3 defines clinically and biologically distinct subgroups of pediatric diffuse intrinsic pontine gliomas. Acta Neuropathol 124: 439–447.

Kloet SL, Makowski MM, Baymaz HI, van Voorthuijsen L, Karemaker ID, Santanach A, Jansen P, Di Croce L, Vermeulen M. 2016. The dynamic interactome and genomic targets of Poly-comb complexes during stem-cell differentiation. Nat Struct Mol Biol 23: 682–690.

Knutson SK, Kawano S, Minoshima Y, Warholic NM, Huang KC, Xiao Y, Kadowaki T, Uesugi M, Kuznetsov G, Kumar N et al. 2014. Selective inhibition of EZH2 by EPZ-6438 leads to potent antitumor activity in EZH2-mutant non-Hodgkin lymphoma. Mol Cancer Ther 13: 842–854.

Knutson SK, Warholic NM, Wigle TJ, Klaus CR, Allain CJ, Raimondi A, Porter Scott M, Chesworth R, Moyer MP, Copeland RA et al. 2013. Durable tumor regression in genetically altered malignant rhabdoid tumors by inhibition of methyltransferase EZH2. Proc Natl Acad Sci U S A 110: 7922–7927.

Krug B, Harutyunyan AS, Deshmukh S, Jabado N. 2021. Poly-comb repressive complex 2 in the driver’s seat of childhood and young adult brain tumours. Trends Cell Biol 31: 814–828.

Kumar SS, Sengupta S, Lee K, Hura N, Fuller C, DeWire M, Stevenson CB, Fouladi M, Drissi R. 2017. BMI-1 is a potential therapeutic target in diffuse intrinsic pontine glioma. Oncotarget 8: 62962–62975.

Lamb KN, Bsteh D, Dishman SN, Moussa HF, Fan H, Stuckey JI, Norris JL, Cholensky SH, Li D, Wang J et al. 2019. Discovery and Characterization of a Cellular Potent Positive Allosteric Modulator of the Polycomb Repressive Complex 1 Chromodomain, CBX7. Cell Chem Biol 26: 1365–1379 e1322.

Langmead B, Salzberg SL. 2012. Fast gapped-read alignment with Bowtie 2. Nat Methods 9: 357–359.

Lerdrup M, Hansen K. 2020. User-Friendly and Interactive Analysis of ChIP-Seq Data Using EaSeq. Methods Mol Biol 2117: 35–63.

Lewis PW, Muller MM, Koletsky MS, Cordero F, Lin S, Banaszynski LA, Garcia BA, Muir TW, Becher OJ, Allis CD. 2013. Inhibition of PRC2 activity by a gain-of-function H3 mutation found in pediatric glioblastoma. Science 340: 857–861.

Li B, Zhou J, Liu P, Hu J, Jin H, Shimono Y, Takahashi M, Xu G. 2007. Polycomb protein Cbx4 promotes SUMO modification of de novo DNA methyltransferase Dnmt3a. Biochem J 405: 369–378.

Li H, Handsaker B, Wysoker A, Fennell T, Ruan J, Homer N, Marth G, Abecasis G, Durbin R, Genome Project Data Processing S. 2009. The Sequence Alignment/Map format and SAM-tools. Bioinformatics 25: 2078–2079.

Li J, Xu Y, Long XD, Wang W, Jiao HK, Mei Z, Yin QQ, Ma LN, Zhou AW, Wang LS et al. 2014. Cbx4 governs HIF-1alpha to potentiate angiogenesis of hepatocellular carcinoma by its SUMO E3 ligase activity. Cancer Cell 25: 118–131.

Liang R, Tomita D, Sasaki Y, Ginn J, Michino M, Huggins DJ, Baxt L, Kargman S, Shahid M, Aso K et al. 2022. A Chemical Strategy toward Novel Brain-Penetrant EZH2 Inhibitors. ACS Med Chem Lett 13: 377–387.

Louis DN, Perry A, Reifenberger G, von Deimling A, Figarel-la-Branger D, Cavenee WK, Ohgaki H, Wiestler OD, Kleihues P, Ellison DW. 2016. The 2016 World Health Organization Classification of Tumors of the Central Nervous System: a summary. Acta Neuropathol 131: 803–820.

Love MI, Huber W, Anders S. 2014. Moderated estimation of fold change and dispersion for RNA-seq data with DESeq2. Genome Biol 15: 550.

Mack SC, Witt H, Piro RM, Gu L, Zuyderduyn S, Stutz AM, Wang X, Gallo M, Garzia L, Zayne K et al. 2014. Epigenomic alterations define lethal CIMP-positive ependymomas of infancy. Nature 506: 445–450.

Mackay A, Burford A, Carvalho D, Izquierdo E, Fazal-Salom J, Taylor KR, Bjerke L, Clarke M, Vinci M, Nandhabalan M et al. 2017. Integrated Molecular Meta-Analysis of 1,000 Pediatric High-Grade and Diffuse Intrinsic Pontine Glioma. Cancer Cell 32: 520–537 e525.

Maertens GN, El Messaoudi-Aubert S, Racek T, Stock JK, Nicholls J, Rodriguez-Niedenfuhr M, Gil J, Peters G. 2009. Several distinct polycomb complexes regulate and co-localize on the INK4a tumor suppressor locus. PLoS One 4: e6380.

Margueron R, Li G, Sarma K, Blais A, Zavadil J, Woodcock CL, Dynlacht BD, Reinberg D. 2008. Ezh1 and Ezh2 maintain repressive chromatin through different mechanisms. Mol Cell 32: 503–518.

Meier F, Brunner AD, Koch S, Koch H, Lubeck M, Krause M, Goedecke N, Decker J, Kosinski T, Park MA et al. 2018. Online Parallel Accumulation-Serial Fragmentation (PASEF) with a Novel Trapped Ion Mobility Mass Spectrometer. Mol Cell Proteomics 17: 2534–2545.

Michealraj KA, Kumar SA, Kim LJY, Cavalli FMG, Przelicki D, Wojcik JB, Delaidelli A, Bajic A, Saulnier O, MacLeod G et al. 2020. Metabolic Regulation of the Epigenome Drives Lethal In-fantile Ependymoma. Cell 181: 1329–1345 e1324.

Min J, Zhang Y, Xu RM. 2003. Structural basis for specific binding of Polycomb chromodomain to histone H3 methylated at Lys 27. Genes Dev 17: 1823–1828.

Mohammad F, Weissmann S, Leblanc B, Pandey DP, Hojfeldt JW, Comet I, Zheng C, Johansen JV, Rapin N, Porse BT et al. 2017. EZH2 is a potential therapeutic target for H3K27M-mutant pediatric gliomas. Nat Med 23: 483–492.

Monje M, Mitra SS, Freret ME, Raveh TB, Kim J, Masek M, Attema JL, Li G, Haddix T, Edwards MS et al. 2011. Hedgehog-responsive candidate cell of origin for diffuse intrinsic pontine glioma. Proc Natl Acad Sci U S A 108: 4453–4458.

Nikbakht H, Panditharatna E, Mikael LG, Li R, Gayden T, Osmond M, Ho CY, Kambhampati M, Hwang EI, Faury D et al. 2016. Spatial and temporal homogeneity of driver mutations in diffuse intrinsic pontine glioma. Nat Commun 7: 11185.

Nikoloski G, Langemeijer SM, Kuiper RP, Knops R, Massop M, Tonnissen ER, van der Heijden A, Scheele TN, Vandenberghe P, de Witte T et al. 2010. Somatic mutations of the histone methyl-transferase gene EZH2 in myelodysplastic syndromes. Nat Genet 42: 665–667.

Ntziachristos P, Tsirigos A, Van Vlierberghe P, Nedjic J, Trimarchi T, Flaherty MS, Ferres-Marco D, da Ros V, Tang Z, Siegle J et al. 2012. Genetic inactivation of the polycomb repressive complex 2 in T cell acute lymphoblastic leukemia. Nat Med 18: 298–301.

Orlando DA, Chen MW, Brown VE, Solanki S, Choi YJ, Olson ER, Fritz CC, Bradner JE, Guenther MG. 2014. Quantitative ChIP-Seq normalization reveals global modulation of the epigenome. Cell Rep 9: 1163–1170.

Pajtler KW, Wen J, Sill M, Lin T, Orisme W, Tang B, Hubner JM, Ramaswamy V, Jia S, Dalton JD et al. 2018. Molecular hetero-geneity and CXorf67 alterations in posterior fossa group A (PFA) ependymomas. Acta Neuropathol 136: 211–226.

Panditharatna E, Marques JG, Wang T, Trissal MC, Liu I, Jiang L, Beck A, Groves A, Dharia NV, Li D et al. 2022. BAF Complex Maintains Glioma Stem Cells in Pediatric H3K27M Glioma. Cancer Discov 12: 2880–2905.

Pemberton H, Anderton E, Patel H, Brookes S, Chandler H, Palermo R, Stock J, Rodriguez-Niedenfuhr M, Racek T, de Breed L et al. 2014. Genome-wide co-localization of Polycomb orthologs and their effects on gene expression in human fibroblasts. Genome Biol 15: R23.

Phillips RE, Yang Y, Smith RC, Thompson BM, Yamasaki T, So-to-Feliciano YM, Funato K, Liang Y, Garcia-Bermudez J, Wang X et al. 2019. Target identification reveals lanosterol synthase as a vulnerability in glioma. Proc Natl Acad Sci U S A 116: 7957–7962.

Piunti A, Hashizume R, Morgan MA, Bartom ET, Horbinski CM, Marshall SA, Rendleman EJ, Ma Q, Takahashi YH, Woodfin AR et al. 2017. Therapeutic targeting of polycomb and BET bromo-domain proteins in diffuse intrinsic pontine gliomas. Nat Med 23: 493–500.

Piunti A, Smith ER, Morgan MAJ, Ugarenko M, Khaltyan N, Helmin KA, Ryan CA, Murray DC, Rickels RA, Yilmaz BD et al. 2019. CATACOMB: An endogenous inducible gene that antagonizes H3K27 methylation activity of Polycomb repressive complex 2 via an H3K27M-like mechanism. Sci Adv 5: eaax2887.

Plys AJ, Davis CP, Kim J, Rizki G, Keenen MM, Marr SK, Kingston RE. 2019. Phase separation of Polycomb-repressive complex 1 is governed by a charged disordered region of CBX2. Genes Dev 33: 799–813.

Quinlan AR, Hall IM. 2010. BEDTools: a flexible suite of utilities for comparing genomic features. Bioinformatics 26: 841–842.

Rappsilber J, Mann M, Ishihama Y. 2007. Protocol for micro-purification, enrichment, pre-fractionation and storage of peptides for proteomics using StageTips. Nat Protoc 2: 1896–1906.

Roscic A, Moller A, Calzado MA, Renner F, Wimmer VC, Gresko E, Ludi KS, Schmitz ML. 2006. Phosphorylation-dependent control of Pc2 SUMO E3 ligase activity by its substrate protein HIPK2. Mol Cell 24: 77–89.

Ryall S, Guzman M, Elbabaa SK, Luu B, Mack SC, Zapotocky M, Taylor MD, Hawkins C, Ramaswamy V. 2017. H3 K27M mutations are extremely rare in posterior fossa group A ependymoma. Childs Nerv Syst 33: 1047–1051.

Sanjana NE, Shalem O, Zhang F. 2014. Improved vectors and genome-wide libraries for CRISPR screening. Nat Methods 11: 783–784.

Scelfo A, Fernandez-Perez D, Tamburri S, Zanotti M, Lavarone E, Soldi M, Bonaldi T, Ferrari KJ, Pasini D. 2019. Functional Landscape of PCGF Proteins Reveals Both RING1A/B-Dependent-and RING1A/B-Independent-Specific Activities. Mol Cell 74: 1037–1052 e1037.

Schwartzentruber J, Korshunov A, Liu XY, Jones DT, Pfaff E, Jacob K, Sturm D, Fontebasso AM, Quang DA, Tonjes M et al. 2012. Driver mutations in histone H3.3 and chromatin remodelling genes in paediatric glioblastoma. Nature 482: 226–231.

Shen C, Ipsaro JJ, Shi J, Milazzo JP, Wang E, Roe JS, Suzuki Y, Pappin DJ, Joshua-Tor L, Vakoc CR. 2015. NSD3-Short Is an Adaptor Protein that Couples BRD4 to the CHD8 Chromatin Remodeler. Mol Cell 60: 847–859.

Shen X, Liu Y, Hsu YJ, Fujiwara Y, Kim J, Mao X, Yuan GC, Orkin SH. 2008. EZH1 mediates methylation on histone H3 lysine 27 and complements EZH2 in maintaining stem cell identity and executing pluripotency. Mol Cell 32: 491–502.

Shi J, Wang E, Milazzo JP, Wang Z, Kinney JB, Vakoc CR. 2015. Discovery of cancer drug targets by CRISPR-Cas9 screening of protein domains. Nat Biotechnol 33: 661–667.

Skene PJ, Henikoff S. 2017. An efficient targeted nuclease strategy for high-resolution mapping of DNA binding sites. Elife 6.

Smits AH, Jansen PW, Poser I, Hyman AA, Vermeulen M. 2013. Stoichiometry of chromatin-associated protein complexes revealed by label-free quantitative mass spectrometry-based proteomics. Nucleic Acids Res 41: e28.

Soto-Feliciano YM, Sanchez-Rivera FJ, Perner F, Barrows DW, Kastenhuber ER, Ho YJ, Carroll T, Xiong Y, Anand D, Soshnev AA et al. 2023. A Molecular Switch between Mammalian MLL Complexes Dictates Response to Menin-MLL Inhibition. Cancer Discov 13: 146–169.

Spahn PN, Bath T, Weiss RJ, Kim J, Esko JD, Lewis NE, Harismendy O. 2017. PinAPL-Py: A comprehensive web-application for the analysis of CRISPR/Cas9 screens. Sci Rep 7: 15854.

Straining R, Eighmy W. 2022. Tazemetostat: EZH2 Inhibitor. J Adv Pract Oncol 13: 158–163.

Suh JL, Bsteh D, Hart B, Si Y, Weaver TM, Pribitzer C, Lau R, Soni S, Ogana H, Rectenwald JM et al. 2022. Reprogramming CBX8-PRC1 function with a positive allosteric modulator. Cell Chem Biol 29: 555–571 e511.

Tarumoto Y, Lu B, Somerville TDD, Huang YH, Milazzo JP, Wu XS, Klingbeil O, El Demerdash O, Shi J, Vakoc CR. 2018. LKB1, Salt-Inducible Kinases, and MEF2C Are Linked Dependencies in Acute Myeloid Leukemia. Mol Cell 69: 1017–1027 e1016.

Tatavosian R, Kent S, Brown K, Yao T, Duc HN, Huynh TN, Zhen CY, Ma B, Wang H, Ren X. 2019. Nuclear condensates of the Polycomb protein chromobox 2 (CBX2) assemble through phase separation. J Biol Chem 294: 1451–1463.

van Mierlo G, Veenstra GJC, Vermeulen M, Marks H. 2019. The Complexity of PRC2 Subcomplexes. Trends Cell Biol 29: 660–671.

Wang B, Wang M, Zhang W, Xiao T, Chen CH, Wu A, Wu F, Traugh N, Wang X, Li Z et al. 2019. Integrative analysis of pooled CRISPR genetic screens using MAGeCKFlute. Nat Protoc 14: 756–780.

Wang L, Brown JL, Cao R, Zhang Y, Kassis JA, Jones RS. 2004. Hierarchical recruitment of polycomb group silencing complexes. Mol Cell 14: 637–646.

Wisniewski JR, Zougman A, Nagaraj N, Mann M. 2009. Universal sample preparation method for proteome analysis. Nat Methods 6: 359–362.

Wu G, Broniscer A, McEachron TA, Lu C, Paugh BS, Becksfort J, Qu C, Ding L, Huether R, Parker M et al. 2012. Somatic histone H3 alterations in pediatric diffuse intrinsic pontine gliomas and non-brainstem glioblastomas. Nat Genet 44: 251–253.

Yang L, Chan AKN, Miyashita K, Delaney CD, Wang X, Li H, Pokharel SP, Li S, Li M, Xu X et al. 2021. High-resolution characterization of gene function using single-cell CRISPR tiling screen. Nat Commun 12: 4063.

Zehir A, Benayed R, Shah RH, Syed A, Middha S, Kim HR, Srinivasan P, Gao J, Chakravarty D, Devlin SM et al. 2017. Mutational landscape of metastatic cancer revealed from prospective clinical sequencing of 10,000 patients. Nat Med 23: 703–713.

Zhang P, de Gooijer MC, Buil LC, Beijnen JH, Li G, van Tellingen O. 2015. ABCB1 and ABCG2 restrict the brain penetration of a panel of novel EZH2-Inhibitors. Int J Cancer 137: 2007–2018.

